# The timing and direction of introgression under the multispecies network coalescent

**DOI:** 10.1101/328575

**Authors:** Mark S. Hibbins, Matthew W. Hahn

## Abstract

Introgression is a pervasive biological process, and many statistical methods have been developed to infer its presence from genomic data. However, many of the consequences and genomic signatures of introgression remain unexplored from a methodological standpoint. Here, we develop a model for the timing and direction of introgression based on the multispecies network coalescent, and from it suggest new approaches for testing introgression hypotheses. We suggest two new statistics, *D*_1_ and *D*_2_, which can be used in conjunction with other information to test hypotheses relating to the timing and direction of introgression, respectively. *D*_1_ may find use in evaluating cases of homoploid hybrid speciation, while *D*_2_ provides a four-taxon test for polarizing introgression. Although analytical expectations for our statistics require a number of assumptions to be met, we show how simulations can be used to test hypotheses about introgression when these assumptions are violated. We apply the *D*_1_ statistic to genomic data from the wild yeast *Saccharomyces paradoxus*, a proposed example of homoploid hybrid speciation, demonstrating its use as a test of this model. These methods provide new and powerful ways to address questions relating to the timing and direction of introgression.

## Introduction

The now-widespread availability of genomic data has demonstrated that gene flow between previously diverged lineages—also known as introgression—is a pervasive process across the tree of life (reviewed in Mallet et al. 2016). Whole-genome data has revealed the sharing of traits via gene flow between humans and an extinct lineage known as Denisovans (Huerta-Sánchez et al. 2014), between different species of *Heliconius* butterflies (The *Heliconius* Genome Consortium 2012), and between multiple malaria vectors in the *Anopheles gambiae* species complex (Fontaine et al. 2015; Wen et al. 2016a). Introgression can substantially alter the evolutionary trajectory of populations through adaptive introgression (Hedrick 2013), transgressive segregation (Rieseberg et al. 1999), and hybrid speciation (Schumer et al. 2014).

Species or populations exchanging migrants are often represented as a network, rather than as having strictly bifurcating relationships. The reticulations in such networks represent the histories of loci that have crossed species boundaries. However, phylogenetic networks are often conceived in different ways (Huson and Bryant 2006). Some representations imply specific evolutionary processes or directions of introgression, but these implications are not always intentional and/or properly addressed. For example, Figure 1 shows three ways in which introgression events can be depicted in a network, with species B and C involved in gene exchange in each. Figure 1a represents an introgression event between species B and C, after A and B have diverged and B has evolved independently for some period of time. This representation does not specify the direction of gene flow. Figure 1b suggests that lineage B is the result of hybridization between A and C, and therefore that the direction of gene flow is into B. Such a depiction is often used to represent the origin of admixed populations (e.g. Bertorelle and Excoffier 1998; Wang 2003), or hybrid speciation (e.g. Meng and Kubatko 2009), in which the hybridization event leads to the formation of a reproductively isolated lineage. Figure 1c suggests two speciation events that result in lineages sister to A and C, respectively, that then come together to form species B. This representation also implies the direction of introgression (into B), but differs from Figure 1b in that it could imply a period of independent evolution before hybridization (e.g. Patterson et al. 2012; Yu et al. 2014; Zhang et al. 2018).

**Figure 1:**
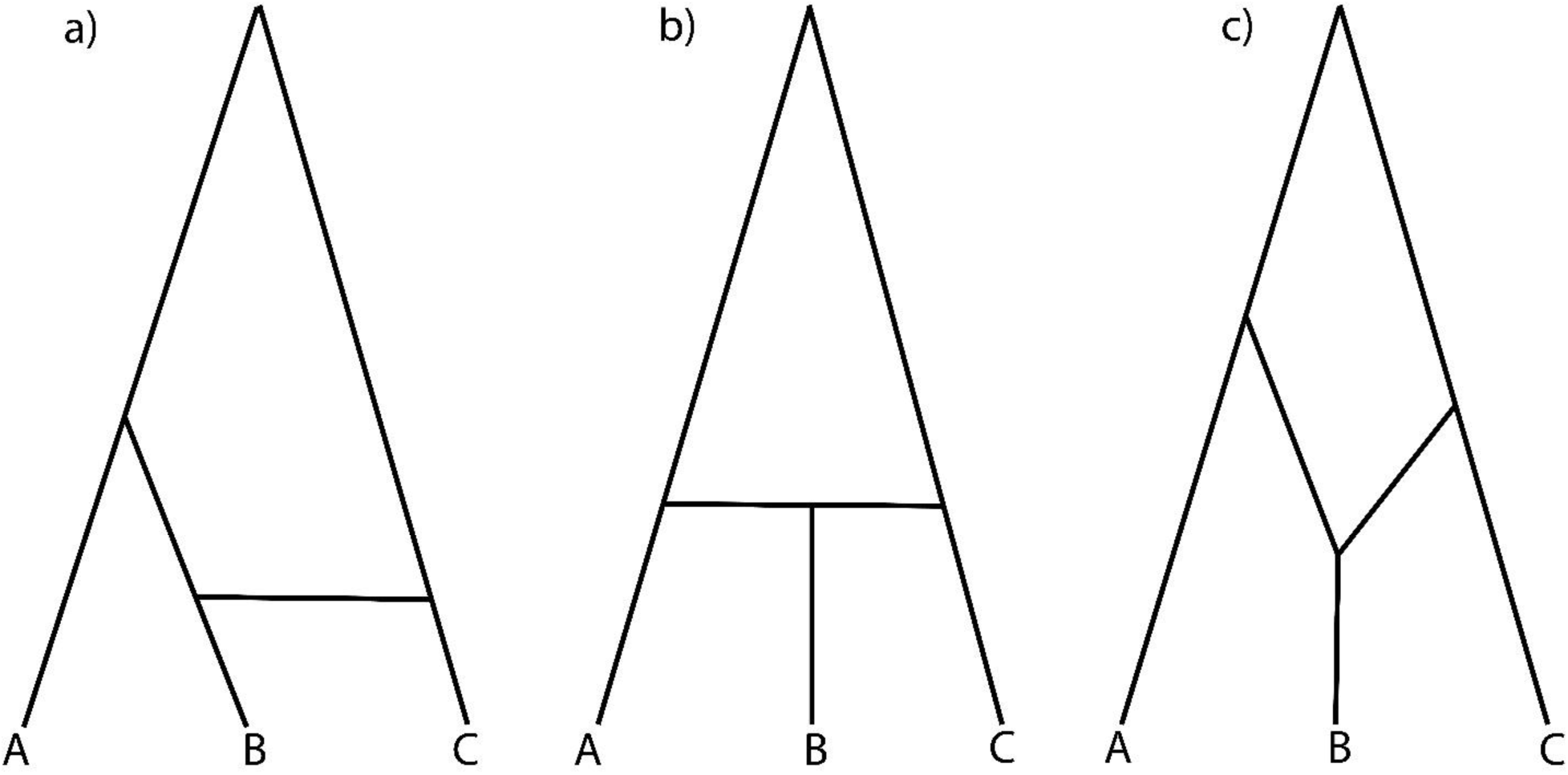
Different phylogenetic network representations of species relationships. a) Speciation between lineages A and B, followed by introgression between lineages B and C (with unspecified direction). b) Homoploid hybrid speciation. Lineage B is created by a hybridization event between A and C. c) This representation is used to denote either speciation between lineages A and B followed by introgression from C into B, or two speciation events followed by the merging of two lineages to form species B.

Despite clearly representing different evolutionary scenarios—including in the timing of introgression relative to speciation, and the direction of introgression—species networks such as these are often used to represent introgression as a general process. One reason for this is that the three scenarios depicted in Figure 1 are difficult to distinguish. Popular methods for detecting introgression using SNPs, such as the *f*_3_ and *f*_4_ statistics (Reich et al. 2009; Patterson et al. 2012) and the related *D* statistic (also known as the “ABBA-BABA” test; Green et al. 2010; Durand et al. 2011) can tell us whether introgression has occurred, and if so, between which taxa. These methods do not provide additional information about the introgression event, including its direction or its relationship to speciation. The same is true of phylogenetic methods that use gene trees without branch lengths to infer phylogenetic networks (e.g. Meng and Kubatko 2009; Yu et al. 2014), as the frequency of discordant topologies are the same under many different scenarios for the timing and direction of gene flow (Zhu and Degnan 2017).

Accurately inferring the direction of introgression and the timing of introgression is important in understanding the consequences of hybridization for evolution. One area where these inferences have become especially important is in evaluating the frequency of homoploid hybrid speciation (HHS). Schumer et al. (2014) proposed three criteria that must be met in order to label a species a homoploid hybrid: evidence for hybridization, evidence for reproductive isolation, and a causal link between the two. In applying these criteria, they suggested that few studies have been able to demonstrate a causal link between hybridization and reproductive isolation, and that HHS is likely a rare process. This has sparked a debate over how to characterize HHS and its frequency in nature (Feliner et al. 2017; Schumer et al. 2018a). Multiple studies of putative homoploid hybrids have tested general hypotheses of gene flow using genomic data, often in combination with morphological and reproductive isolation data (e.g. Elgvin et al. 2017; Barrera-Guzman et al. 2018). However, to date no population genetic models have been formulated that can provide explicit quantitative predictions of HHS. Such predictions would provide invaluable in characterizing the prevalence, causes, and consequences of HHS.

To address the aforementioned problems, here we develop an explicit model for the timing and direction of introgression based on the multispecies network coalescent (Yu et al. 2012; Yu et al. 2014; Wen et al. 2016b). The multispecies network coalescent model generalizes the multispecies coalescent (Hudson 1983; Rannala and Yang 2003) to allow for both incomplete lineage sorting and introgression (reviewed in Degnan 2018; Elworth et al. 2018). Under this model, a single sample taken from each of the extant lineages traces its history back through the network, following alternative paths produced by reticulations with a probability proportional to the amount of introgression that has occurred. The multispecies network coalescent model differs from population genetic approaches requiring multiple samples per population (such as the “IM” model; Nielsen and Wakeley 2001), though it does require that at least three species (and usually an outgroup) are sampled. However, the use of more than two lineages also makes it possible to more finely resolve the timing of migration, a problem that exists when analyzing sister species in the IM framework (Sousa et al. 2011).

Our study provides expectations for pairwise coalescence times under the multispecies network coalescent model. Two new statistics arise from these expectations, dubbed *D*_1_ and *D*_2_, which can be used alongside other information to test hypotheses regarding the timing and direction of intogression. We perform simulations to establish the power of these statistics when speciation and hybridization have occurred at various times in the recent past, with varying admixture proportions and effective population sizes. Finally, in order to demonstrate the use of the *D*_1_ statistic, we apply it to a genomic dataset from the wild yeast species *Saccharomyces paradoxus*, a potential case of homoploid hybrid speciation (Leducq et al. 2016).

## Materials and Methods

### A multispecies network coalescent model of introgression

Many statistics for detecting introgression, including the ABBA-BABA test, are based on expectations for the frequencies of different gene tree topologies. Our model, and the statistics that follow from it, are instead based on expected coalescence times—and the resulting levels of divergence—between pairs of populations or species. In what follows we present explicit expressions for these coalescence times, for genealogies evolving within a species network. Later in the paper we show how these times can complement and extend analyses based solely on the frequency of different topologies.

To make it easier to track the history of individual loci, we imagine that a species network can be separated into one or more “parent trees” (cf. Meng and Kubatko 2009; Liu et al. 2014). Every reticulation event in the network produces another parent tree, and loci with a history that follows such reticulate branches are considered to be produced by the corresponding parent tree. Embedded in a species network like the one shown in Figure 1a (where introgression is instantaneous) is one parent tree that represents the speciation history of lineages, which we define more formally here. Consider three taxa that have the phylogenetic relationship ((A,B),C). Let *t*_1_ denote the time of speciation between A and B, and let *t*_2_ denote the time of speciation between the common ancestor of A and B and lineage C (Figure 2). While speciation in nature is not an instantaneous process, in this idealized model these times represent the average split across loci. We additionally define *N* as the effective population size of the ancestral population of all three taxa (i.e. the ancestor of taxa A, B, and C). These relationships, which depict the speciation history of the clade, will be referred to as “parent tree 1” (Figure 2a).

**Figure 2:**
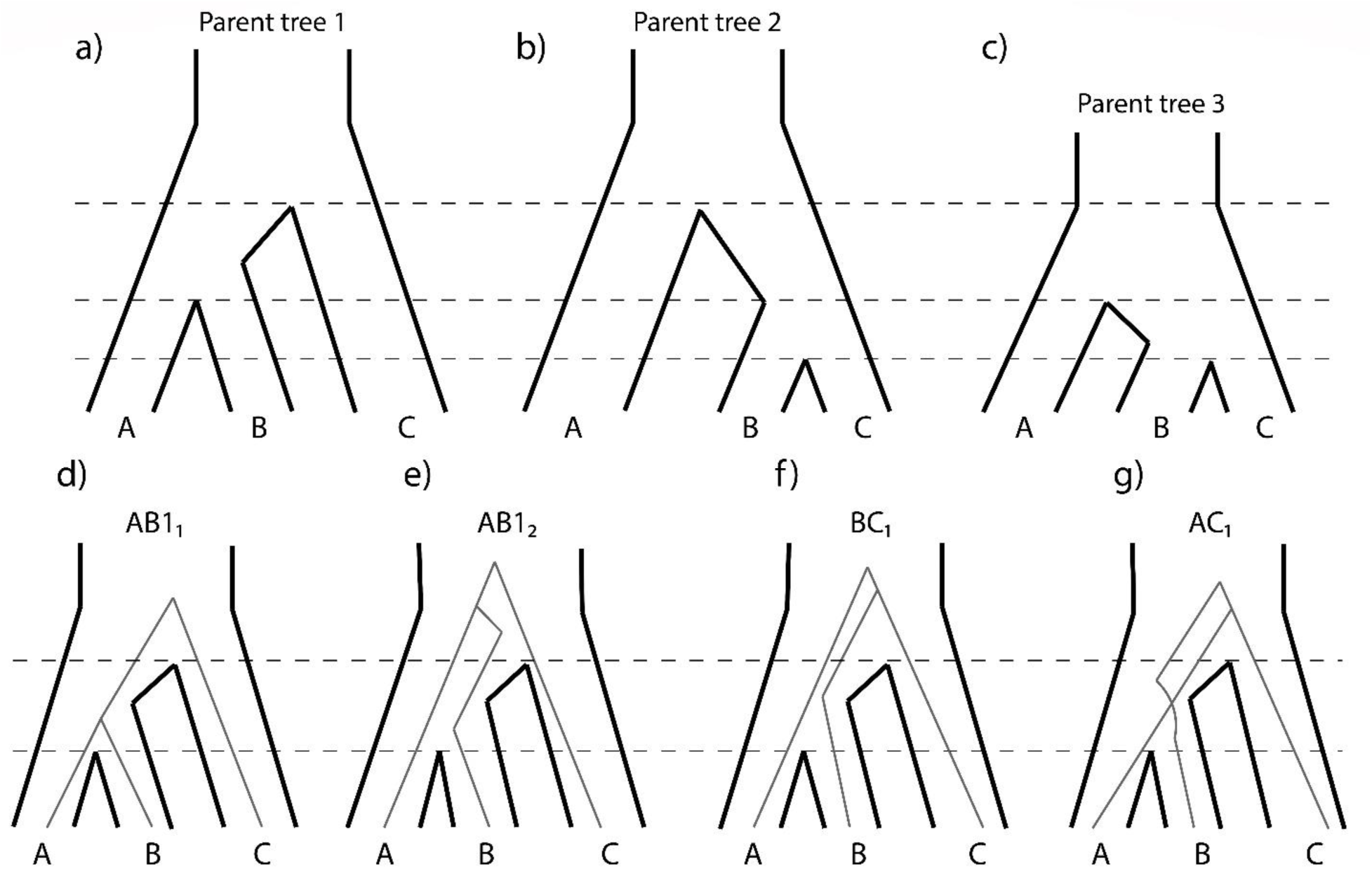
Trees forming the conceptual foundations of our coalescent model, labelled with relevant time parameters. Panels a-c depict the parent trees generated from different introgression scenarios, within which gene trees will sort at particular loci. Panels d-g show the possible ways that gene trees can sort within parent tree 1, demonstrating how gene trees are expected to sort within their respective parent trees according to the coalescent process.

Because of the stochasticity of the coalescent process, parent tree 1 will generate one of four topologies at a particular locus: 1) A concordant tree in which A and B coalesce before *t*_2_, denoted *AB1*_*1*_ (Figure 2d); 2) A concordant tree in which A and B coalesce after *t*_2_, denoted *AB2*_*1*_ (Figure 2e); 3) A discordant tree where B and C are the first to coalesce after *t*_2_, denoted *BC*_*1*_ (Figure 2f); and 4) A discordant tree where A and C are the first to coalesce after *t*_2_, denoted *AC*_*1*_ (Figure 2g). The expected frequency of each of these topologies is a classic result from coalescent theory (Hudson 1983; Tajima 1983; Pamilo and Nei 1988). The expected coalescence times for each pair of species in each topology can also be found using straightforward properties of the coalescent model. There are three possible pairs of species, and therefore three times to coalescence in each of the four topologies. Fortunately, the symmetry of relationships means that many of these times are the same between pairs of species and across topologies. The equations that follow are presented in units of 2*N* generations for simplicity; see the Supplementary Materials for the full expressions.

For the gene tree *AB1*_*1*_ in parent tree 1, the expected time to coalescence between A and B (*t*_A-B_) is:

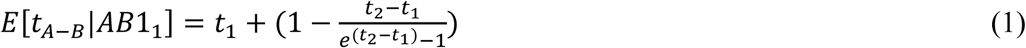

(equation A.7 in Mendes and Hahn 2018). Implicit in this equation is the effective population size of the internal branch of parent tree 1 (i.e. the ancestor of taxa A and B), which determines the length of the branch in coalescent units; from here forward, this population size is referred to as *N*_1_. For the time to coalescence between pairs B-C and A-C in topology *AB1*_*1*_, the lineages must coalesce after *t*_2_ (looking backward in time), and then the lineage ancestral to A and B is expected to coalesce with C in 2*N* generations. Therefore

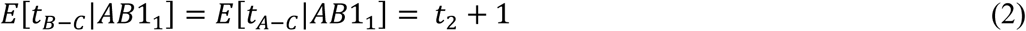

For topology *AB2*_*1*_, A and B now coalesce after *t*_2_, but before either coalesces with another lineage. The time to coalescence is therefore:

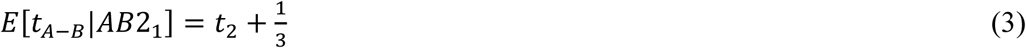

For pairs B-C and A-C in topology *AB2*_*1*_, first A and B must coalesce (which takes 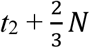 generations on average), and then the lineage ancestral to A and B can coalesce with C. This means that the total expected time for both pairs is:

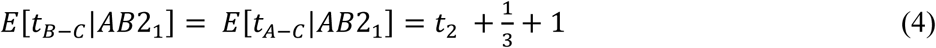

In the discordant topology *BC*_*1*_, species B and C coalesce before any other pair of taxa, and must do so in the ancestral population of all three lineages (after *t*_2_). The time to coalescence is therefore the same as in equation 3:

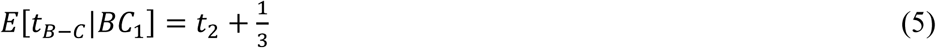

Similarly, the time to the common ancestor of pairs A-B and A-C in topology *BC*_*1*_ follow the same coalescent history as the two pairs in equation 4:

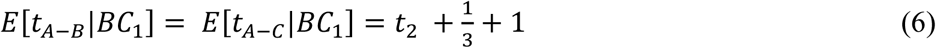

Finally, we have the topology *AC*_*1*_, in which species A and C coalesce first. Their time to coalescence is:

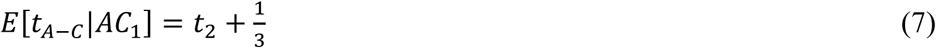

Likewise, the times to the common ancestor of pairs A-B and B-C both take the whole height of the tree to coalesce, so:

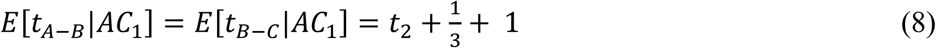

The above expectations were derived under “parent tree 1,” which represents the species tree. If there is introgression from species C into species B, individual loci with a history of introgression will now follow an alternative route through the species network, and therefore a new parent tree. Given that an introgression event between species B and C occurs before *t*_1_ (looking back in time), the topology of this parent tree will be ((B,C),A). We refer to this topology as “parent tree 2” (Figure 2b). The expected coalescence times of gene trees evolving inside parent tree 2 are similar to those from parent tree 1, with a few key differences. First, because this parent tree has the topology ((B,C),A), the two “concordant” gene trees also have this topology. In other words, the coalescent expectations for species pair A-B in parent tree 1 are the same as those for pair B–C in parent tree 2. Second, although *t*_2_ remains the same in the two trees, the first lineage-splitting event is determined by the timing of the instantaneous introgression event in parent tree 2—which we will denote as *t*_m_—rather than by the speciation time, *t*_1_ (Figure 2b). Lastly, parent tree 2 can have a different internal-branch effective population size, which we here denote as *N*_2_.

With these differences in mind, the expected coalescence times for each pair of species in each gene tree topology evolving in parent tree 2 are as follows:

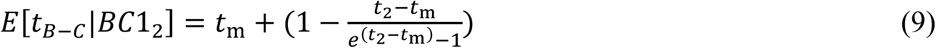

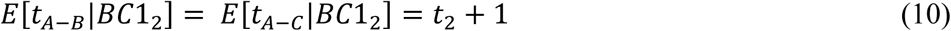

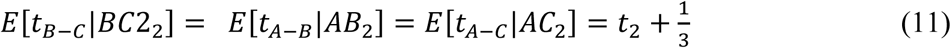

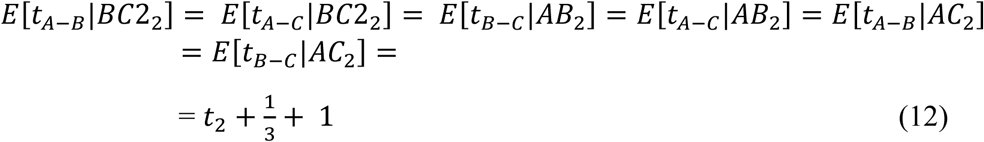

Many parent trees can be defined within a species network, based on the direction of introgression and the taxa involved in introgression. Here we consider one additional tree, denoted parent tree 3 (Figure 2c), which again represents gene flow between species B and C, but in this case from B into C. In this parent tree, the speciation time between species A and B remains *t*_1_, but now this time also represents the first point at which A and C can coalesce—*t*_2_ is not relevant. This is because the presence of loci from lineage B in lineage C allows C to trace its ancestry through B going back in time, which in turn allows it to coalesce with A after *t*_1_. The time to first coalescence for lineages from B and C inside this tree is again limited by the timing of introgression, which again predates *t*_1_, and which we assume occurs at the same time as introgression in parent tree 2 (i.e. *t*_m_). We define the internal-branch effective population size for this parent tree as *N*_3_.

In general, then, the difference between parent tree 3 and parent tree 2 is that all *t*_2_ terms are replaced with *t*_1_. Therefore:

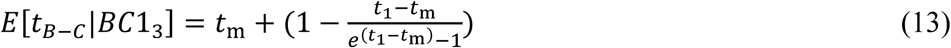

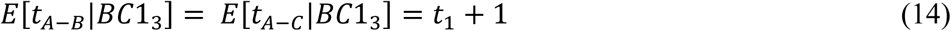

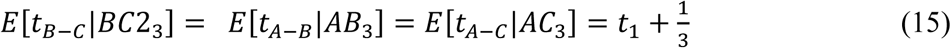

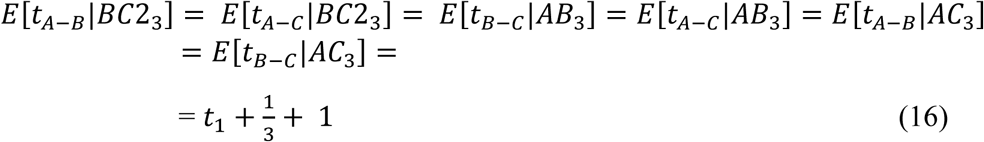

### The *D*_1_ statistic for the relative timing of introgression

Given the expectations laid out above, we can now develop statistics that differentiate alternative biological scenarios. The first comparison we wish to make is between models of speciation followed by introgression (Figure 1a) and models where speciation and introgression are simultaneous (Figure 1b); the latter scenario corresponds to homoploid hybrid speciation or the creation of a new admixed population. We assume for now that introgression has occurred in the direction from C into B in both scenarios, as such cases will be the hardest to distinguish.

The distinguishing feature between these two biological scenarios is the timing of introgression relative to speciation or lineage-splitting, and therefore the expected coalescence times between sequences from species B and either species A or C. If introgression occurs after speciation, we expect that loci which follow the species tree embedded in the network (i.e. parent tree 1) will coalesce further back in time than loci that follow the introgression history in the network (i.e. parent tree 2). Figure 3 demonstrates this expected difference graphically. Explicitly, the expected coalescent times between A and B from parent tree 1 largely mirror the times between B and C from parent tree 2, except for genealogies that coalesce before *t*_2_, where *E*[*t*_*A−B*_|*AB*1_1_] depends on *t*_1_ (equation 1), while *E*[*t*_*B−C*_|*BC*1_2_] depends on *t*_m_ (equation 9).

**Figure 3:**
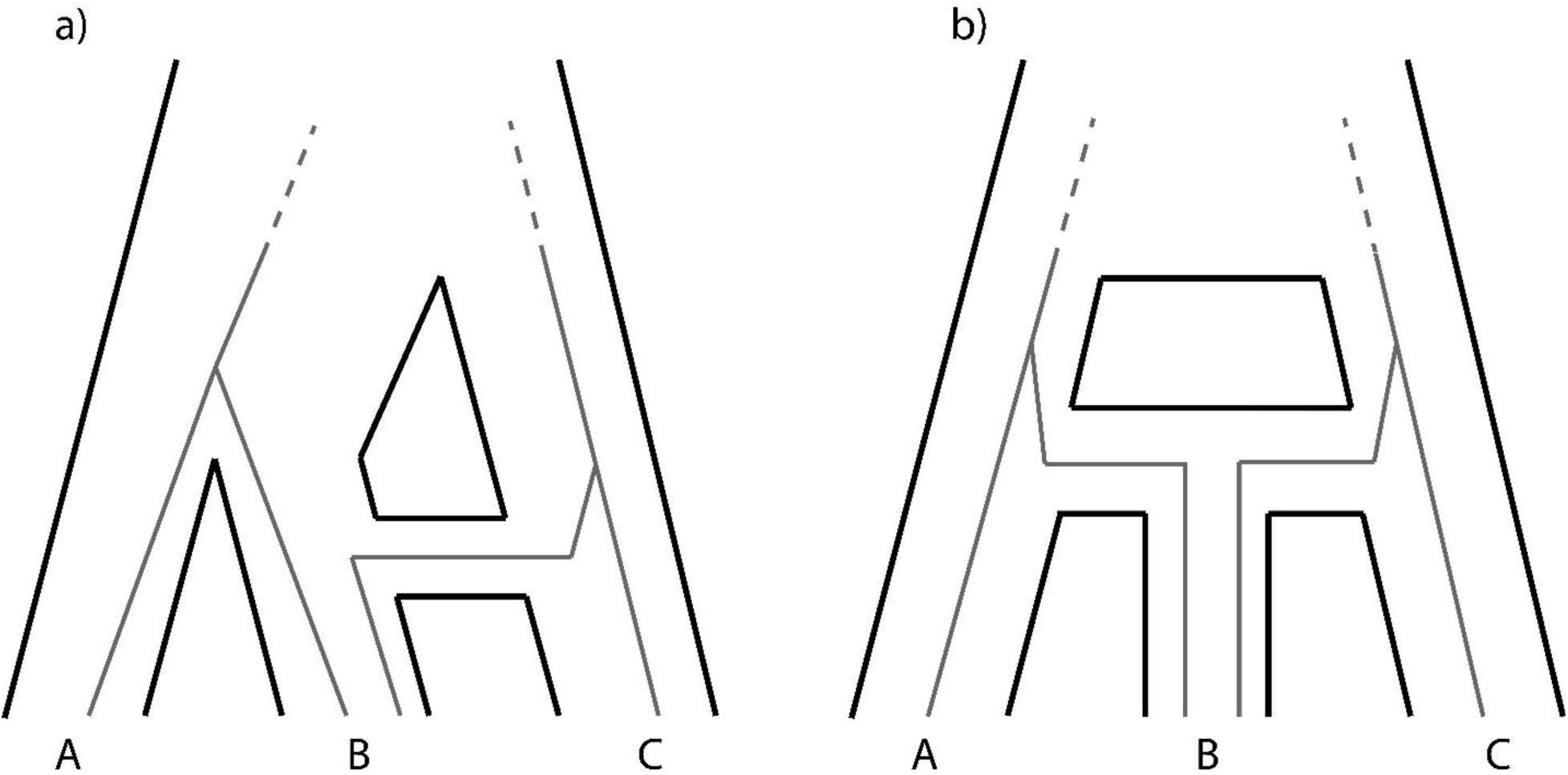
Coalescent times depend on the underlying reticulation history. Grey lines show example genealogical histories (for both the *AB* and *BC* topologies), focusing on the timing of the first coalescent event. a) A reticulation history where speciation and introgression do not occur at the same time. b) A reticulation history where speciation and introgression occur at the same time (as in homoploid hybrid speciation).

This difference captures information on the timing of introgression relative to speciation. If *t*_1_ and *t*_m_ are equal, speciation and introgression are effectively simultaneous, as would be the case in homoploid hybrid speciation. In this case, *E*[*t*_*A−B*_|*AB*1_1_] = *E*[*t*_*B−C*_|*BC*1_2_], and the expected difference between these times is 0. If introgression has occurred significantly before speciation (going backward in time), *t*_1_ > *t*_m_, and therefore, *E*[*t*_*A−B*_|*AB*1_1_] − *E*[*t*_*B−C*_|*BC*1_2_] > 0. This difference will be larger the more recent the introgression event is relative to speciation.

Importantly, this expectation is conditional on *N*_1_ and *N*_2_ being equal; if they are not, the patterns produced by the alternative scenarios will depend both on the timing of introgression relative to speciation and the degree and direction of variation in *N*. Changing the value of *N*_2_ affects the rate of coalescence after time *t*_m_ and the height of genealogies coming from parent tree 2. However, the model presented in the previous section can incorporate this variation, and it can be incorporated into our test statistic (see below).

In developing a statistic to distinguish these scenarios that can be applied to real data, there are two important things to note. First, expected coalescent times can easily be used to model expected amounts of divergence through a simple multiplication by *2μ*, where *μ* is the mutation rate per generation (assuming a constant mutation rate throughout the tree). As divergence can be measured directly from sequence data, the statistics presented here will be in terms of divergence. Second, we cannot know whether the gene tree topology at any given locus was generated by introgression or by incomplete lineage sorting. More importantly, we also do not know if the gene tree originates from parent tree 1 or parent tree 2. This affects both the theoretical expectation and empirical calculation of any statistic.

Ideally, in our formulation of a test statistic, we would like to ignore all irrelevant terms and simply take the difference *E*[*t_A−B_*|*AB*1_1_] − *E*[*t_B−C_*|*BC*1_2_]. However, due to the aforementioned practical constraints, this is not possible. In a dataset where the only available data for a given locus is the gene tree topology and pairwise genetic divergence, the most useful test would measure genetic divergence conditional on a specific tree topology. While we cannot assign parent trees to each gene tree, we know from the coalescent process that the majority of gene trees with the topology ((A,B),C) will come from parent tree 1, and likewise that the majority of gene trees with the topology ((B,C),A) will come from parent tree 2. Therefore, if we measure the distance between A and B in all trees with the ((A,B),C) topology (denoted *d*_A−B_|*AB*), and the distance between B and C in all trees with the ((B,C),A) topology (*d*_B−C_|*BC*), we should be able to capture most of the difference in coalescence times caused by differences between *t*_1_ and *t*_m_.

Based on these considerations, we define a statistic to test the hypothesis that *t*_1_=*t*_m_ as:

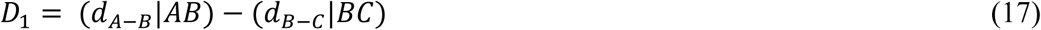

In terms of the coalescence times and genealogies defined above, weighting each expectation by the frequency of the relevant gene tree (denoted with *f* terms):

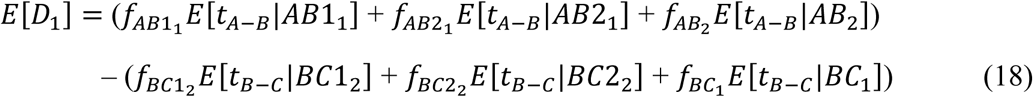

These expectations explicitly recognize that the origin of any genealogy cannot be known, so that *D*_1_ must average across all loci with the same genealogy (either *AB* or *BC*). Expectations for the frequencies of gene trees are given in the Supplementary Materials.

This definition of the *D*_1_ statistic introduces another factor that adds complexity to the expected patterns of divergence: the admixture proportion, which we denote as γ. We call γ_2_ the fraction of gene trees following the introgression history through the network (i.e. originating from parent tree 2). This parameter therefore determines the fraction of gene trees that come from parent tree 1 as well; see the Supplementary Materials for equations describing these expectations explicitly. The effect of γ on gene tree frequencies also interacts with variation in *N*; while γ describes the fraction of loci that originate from a particular parent tree, *N* determines the distribution of topologies that such a parent tree is expected to generate. Under the assumptions that *N*_1_ = *N*_2_ and γ_2_ = 0.5, we expect *D*_1_ to take on a value of 0 for the case of hybrid speciation and a positive value for the case of introgression after speciation. In all other circumstances, the expected value of *D*_1_ under hybrid speciation will depend on the interplay of *N*_1_, *N*_2_ and γ_2_ (see equation SXX in Supplementary Materials).

### The *D*_2_ statistic for the direction of introgression

We also wish to make the distinction between different directions of introgression in terms of our model. As explained above, introgression from C into B generates a different reticulation in the species network, and therefore a different parent tree, than introgression from B into C (compare Figure 2b to 2c). Gene flow from lineage B into lineage C allows loci sampled from C currently to trace their history back through B, which in turn allows lineages A and C to coalesce more quickly (specifically, after *t*_1_ instead of after *t*_2_). Conversely, gene flow from lineage C into lineage B does not change our expectations for the coalescence time of A and C relative to those expected from the species relationships. We can take advantage of these expected differences in the amount of divergence between A and C to develop a statistic for inferring the direction of introgression.

When introgression occurs in the C into B direction, only parent trees 1 and 2 are relevant. The coalescent expectations between A and C from parent tree 1 (equations 2, 4, 6, and 7) exactly mirror those from parent tree 2 (equations 10, 11 and 12). When introgression occurs from B into C, parent trees 1 and 3 become relevant. Coalescence times between A and C in parent tree 3 are all truncated, depending on *t*_1_ instead of *t*_2_ (equations 14, 15 and 16). An obvious distinction between histories is therefore the distance between A and C. In a manner similar to how we defined *D*_1_, we can measure divergence between A and C conditional on alternative gene trees—either ((A,B),C) or ((B,C),A)—with the expectation that the former topology will arise primarily from parent tree 1, and the latter primarily from parent tree 2 or 3. Therefore, we define our statistic for inferring the direction of introgression as:

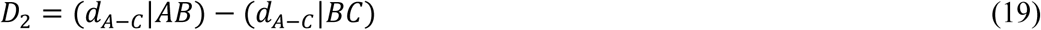

The expected value of *D*_2_ will depend on the direction in which gene flow has occurred. In the case of introgression from C into B, using the same formulation as for *D*_1_, we have:

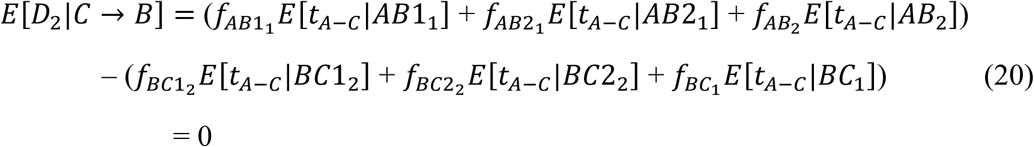

This expression represents the expectation of *D*_2_ when all of our assumptions have been met: when introgression has occurred only from C into B, the statistic should not be significantly different from 0 because the distance between A and C is the same in parent trees 1 and 2. When introgression has occurred from B into C, we have:

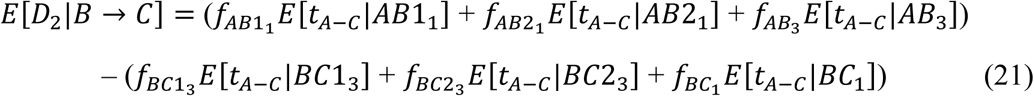

In the expectation of our *D*_2_ statistic, all the relevant gene trees coalesce after *t*_2_, and therefore their rates of coalescence are determined by the same *N*. Because of this, we expect variation in *N* to be less relevant to *D*_2_. However, unequal ancestral population sizes will still affect the expectation of this statistic for introgression from B into C, and the expectations for both directions are expected to be affected by γ_2_ or γ_3_ (see Supplementary Materials). Therefore, like the *D*_1_ statistic, we expect the null hypothesis to be 0 and the alternative to be a positive value under the assumptions of equal population sizes and γ_2_ or γ_3_ = 0.5. In other cases, the expected values of the statistics will come from an interplay of these parameters (see Supplementary Materials).

### Simulations

To investigate the behavior of *D*_1_ and *D*_2_, we explored the parameter space of our model using simulated genealogies from the program *ms* (Hudson 2002). To simulate an introgression event, we took two different approaches in order to ensure the robustness of our results. In the first approach, gene trees were simulated from parent trees separately and then combined in a single dataset, with the *t*_m_ parameter specified for parent tree 2 or 3 representing the timing of the introgression event. In the second approach, we simulated gene trees directly from a species network by specifying an effectively instantaneous population splitting and merging event from the donor to the recipient at time *t*_m_. The specific command lines used in *ms* for both approaches can be found in the Supplementary Materials.

For each statistic, we investigated the effects of the parameters defining the null and alternative hypotheses; for *D*_1_ this is the difference in the timing of speciation and introgression (*t*_1_ − *t*_m_), and for *D*_2_ it is the difference in the timing of lineage-splitting events (*t*_2_ − *t*_1_). Values of the statistics were calculated directly from the branch lengths of simulated genealogies. We also investigated the effects of variation in the effective population size (specifically, the value of *N*_1_ relative to *N*_2_ or *N*_3_) and admixture proportion (γ_2_ or γ_3_). Lastly, we investigated the effect that introgression in both directions has on our statistics. We performed 100 simulations of 2000 genealogies each, for seven different values each of *t*_1_ − *t*_m_, *t*_2_ − *t*_1_, *N*_1_ / *N*_2_ or *N*_1_ / *N*_3_, and γ_2_ or γ_3_, unless specified otherwise. The relevant parameters were simulated for each direction for the *D*2 statistic. We also performed a set of simulations for both statistics in which introgression occurred in both directions rather than only one.

We used a parametric bootstrap approach to evaluate the statistical significance of simulated *D* statistics. We used a two-tailed significance test in which the observed value of the statistic is given a rank *i* in relation to the simulated null distribution, and the *P*-value is calculated as:

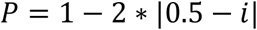

(Voight et al. 2005). We used the same approach to estimate the false negative rate and false positive rate for each combination of parameters. These values were, respectively, the proportion of simulated statistics that incorrectly accept or reject the null hypothesis at a particular significance level.

### Data from *Saccharomyces paradoxus*

To demonstrate the use of our model to test for homoploid hybrid speciation, we analyzed genomic data from three lineages (and an outgroup) of the North American wild yeast species *Saccharomyces paradoxus* (Leducq et al. 2016). This study identified three genetically distinct populations of *S. paradoxus*: two parent lineages dubbed *SpB* and *SpC*, and a hybrid lineage dubbed *SpC*\*. Analysis of whole-genome sequences from these populations shows that *SpC*\* has a mosaic genome, the majority of which is similar to *SpC*, with small genomic regions that are more similar to *SpB*. An investigation of reproductive isolation among all three lineages led Leducq et al. (2016) to conclude that *SpC*\* represents a case of homoploid hybrid speciation.

To evaluate this hypothesis, genomic data from all 161 strains was acquired from the authors. We then separated aligned genomic 5-kb windows into two categories, depending on the assigned topology in Leducq et al. (2016). Windows assigned “ANC” by Leducq et al. represent loci where the topology has *SpC*\* sister to *SpC*; in terms of our definition of *D*1, the distance between *SpC*\* and *SpC* at these loci corresponds to *d_A−B_*|*AB*, under the assumption that the alternative history to hybrid speciation has *SpC*\* and *SpC* as sister species. There are a number of genomic windows in which the topology has *SpC*\* and *SpB* as sister lineages, but some of them are only found in particular groups of strains. We picked windows assigned as “H0” (found in all strains) and “H1b” (found in all strains but one) by Leducq et al. (2016) for our analysis. The distance between *SpC*\* and *SpB* in these windows corresponds to *d*_*B−C*_|*BC* in our formulation of *D*_1_.

For all 161 strains in each of the three lineages (and an outgroup), the total dataset consists of sequence alignments from ANC-topology windows and H0/H1b-topology windows. To carry out calculations of *D*_1_ we then randomly chose windows from 100 different combinations of a single strain from each of *SpC*\*, *SpC* and *SpB*. For each of these 100 samples, we calculated *d_A−B_*|*AB* and *d_B−C_*|*BC* in 5-kb windows simply as the proportion of nucleotide sites that differed between strains; levels of divergence in this system are low enough that no correction for multiple hits is necessary (see Results). For the ANC-topology windows, we calculated *d_A−B_*|*AB* in every other window to reduce the effects of autocorrelation between windows in close physical proximity; the H0/H1b-topology windows were sufficiently few in number and spaced far enough apart so that this was unnecessary for *d_B−C_*|*BC*. *D*_1_ was calculated as the difference in the mean of these two groups. Filtering and distance calculations from all genomic windows were carried out using the software package MVFtools (Pease and Rosenzweig 2018).

To evaluate our observed distribution of *D*_1_ statistics with respect to the HHS hypothesis, we performed a set of simulations corresponding to an HHS scenario using parameters estimated from the study. First, we calculated average genome-wide per-site expected heterozygosity (π) as an estimator for the population-scaled mutation rate, *θ*, in both *SpC* and *SpB*. We used the estimates of π from these populations as proxies for *θ* along the internal branches of the ANC and H0/H1b topologies, respectively. To estimate the population parameters *N*_1_, *N*_2_, *t*_2_, and *t*_1_, we used our estimates of *θ* in conjunction with per-generation mutation rates and generation times from *S. cerevisiae* (Fay and Benavides 2005, Zhu et al. 2014), as well as divergence time estimates from Leducq et al. (2016). We simulated 10000 datasets under the assumption that *t*_1_ and *t*_m_ are equal (as would be the case under HHS). Each dataset consisted of 2002 “ANC” loci and 55 “H0/H1b” loci, sampled from the relevant parent tree.

### Data availability

Code used to generate the simulated genealogical data is available in the Supplementary Materials. Genomic alignments used in the analysis of *S. paradoxus* are from Leducq et al. (2016), and are uploaded to figshare through the GSA portal at https://gsajournals.figshare.com/submit along with the Supplementary Materials.

## Results

### Power of *D*_1_ to distinguish alternative histories

To determine the power of our new statistic, *D*_1_, we asked whether it could distinguish between a history of speciation followed by introgression and a history of homoploid hybrid speciation (HHS), under different regions of the parameter space of our model. Under HHS, speciation and introgression happen simultaneously (i.e. *t*_1_=*t*_m_), and the expected value of *D*_1_ is 0 (equations 17 and 18), assuming that *N*_1_ = *N*_2_ and γ_2_ = 0.5. As the time between speciation and introgression increases, the value of *D*_1_ should increase linearly.

As expected from our model, simulated values of *D*_1_ are centered around a mean of 0 when *t*_1_*−*t_m_=0, *N*_1_ = *N*_2_, and γ_2_ = 0.5, with values of *D*_1_ increasing linearly as the introgression event becomes more recent relative to speciation (Figure 4a). In this region of parameter space introgression that occurs shortly after speciation may be difficult to distinguish from HHS; we observe false negative rates (incorrectly accepting null hypothesis of HHS) of 58 and 70% when *t*_1_ − *t*_m_ = 0.05 (Supplementary Table S2). Values of *t*_1_ − *t*_m_ of 0.1 or greater can always be distinguished from HHS under these assumptions (Supplementary Table S2).

One factor that may reduce the power of *D*_1_ is the presence of introgressed topologies from additional parent trees. Our model predicts that *D*_1_ should have reduced power when introgression occurs in both directions (i.e. C→B and B→C), due to the presence of gene trees generated by parent tree 3. The most common gene trees generated by parent tree 3 have the same topology as those generated by parent tree 2, but the shorter coalescence time between lineages A and B in this parent tree make tests based on *D*_1_ conservative. To investigate the magnitude of this effect, we simulated across the same values of *t*_1_−*t*_m_, now specifying equal contributions of parent trees 2 and 3. Figure 4b shows that introgression shortly after speciation in both directions results in negative values of *D*_1_, and that *D*_1_ can be centered around 0 when *t*_1_− *t*_m_ is not 0. Here, the relative magnitude of γ_2_ and γ_3_ matters: increasingly large values of γ_3_ relative to γ_2_ will magnify this effect.

We expected two other parameters to affect the value of *D*_1_: the difference in effective population size (*N*_1_/*N*_2_) and the admixture proportion (γ_2_). Differences in *N* affect the coalescence times of individual genealogies as well as the degree of incomplete lineage sorting within each parent tree, while the admixture proportion interacts with the degree of incomplete lineage sorting to determine the likely parent tree of origin for a particular gene tree topology. To investigate these effects, we simulated across seven values both of *N*_1_/*N*_2_ and γ_2_. Our results (Figure 5; Supplementary Tables S1 and S2) show that the *D*_1_ statistic is affected by variation in both *N*_1_/*N*_2_ and γ_2_. A two-fold difference in *N* or a difference of 15% in γ_2_ are enough to essentially guarantee that, in the case of HHS, *D*_1_ will deviate significantly from the idealized expectation of 0. There are also regions of parameter space in which the statistic is not likely to deviate from 0, even when there has been introgression after speciation. This occurs when the effect of either *N*_1_ / *N*_2_ or γ_2_ effectively “cancels out” the signals of divergence, leading to *D*_1_ values close to 0. This risk appears to be highest when *N* is smaller in parent tree 1 (Figure 5a), or for values of γ_2_ between 0.05 and 0.5 (Figure 5b). Our expectations for *D*_1_ can also take into account variation in *N* and γ, if these quantities can be estimated (Supplementary equations 1 – 20). In such cases—and when simulations are used to generate the null (see below)—we do not have to use the expectation that *D*_1_ = 0.

**Figure 4:**
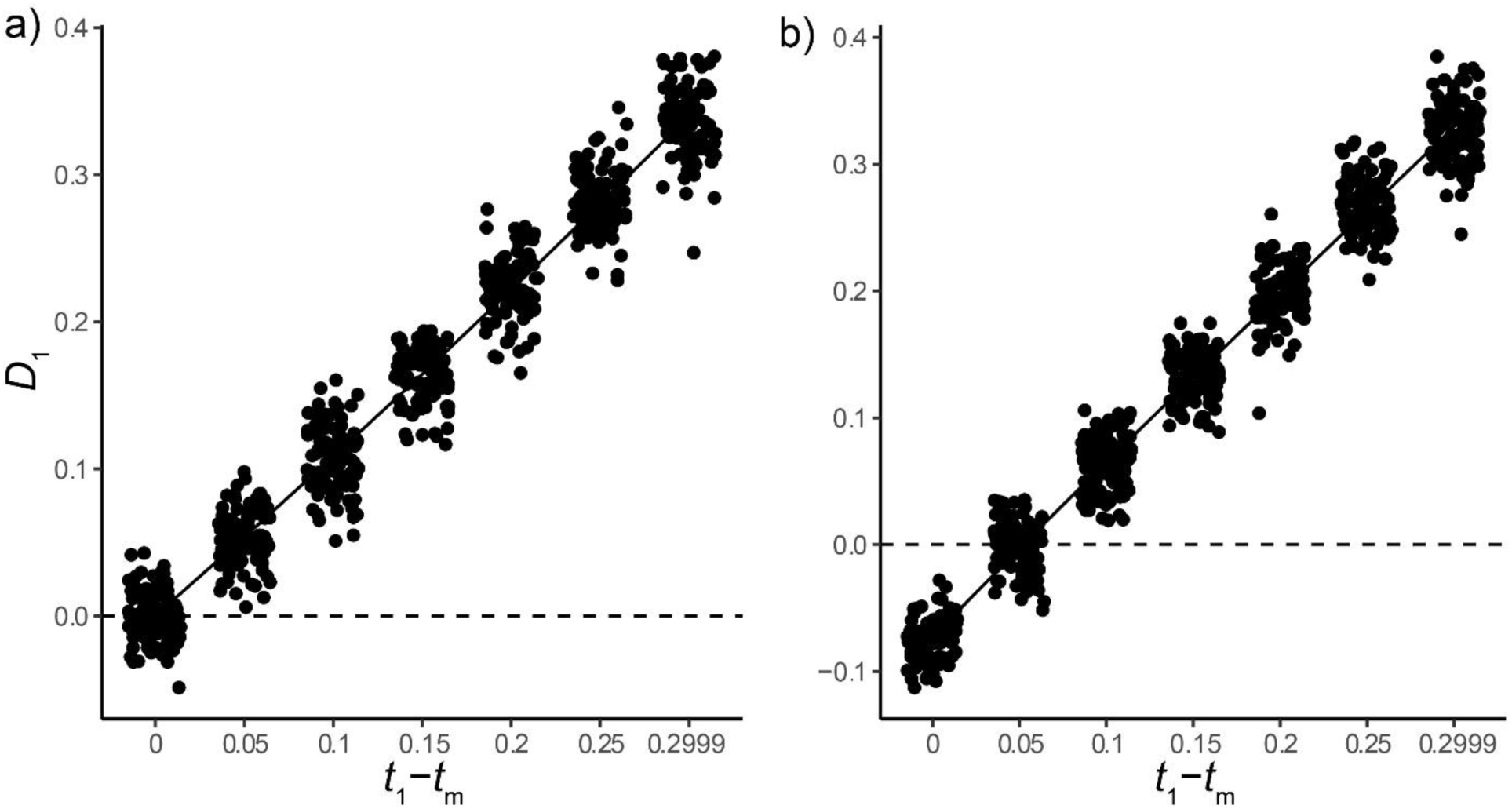
*D*_1_ as a function of the difference in the timing of speciation and introgression, for a) introgression in the C→B direction only, and b) introgression in both directions. Dots represent values obtained from simulations, with jitter added for clarity. The solid line shows the expected values from our coalescent model, while the dashed line shows the null hypothesis of *t*_1_ − *t*m = 0. Time on the x-axis is measured in units of 4*N*.

**Figure 5:**
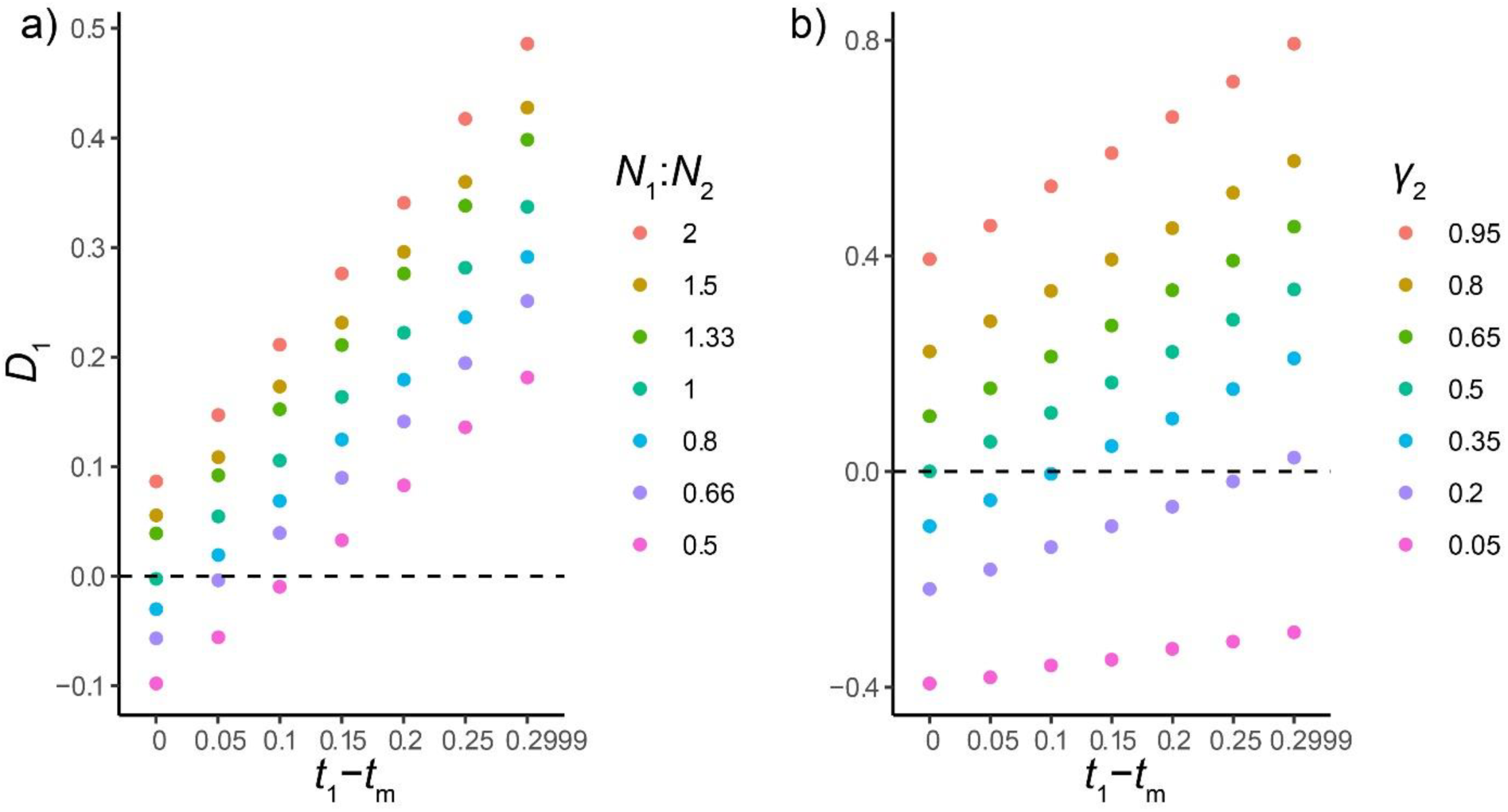
*D*_1_ as a function of variation in *N*_1_ / *N*_2_ and γ_2_. a) Simulations show that variation in *N*_1_/ *N*_2_ (color legend) changes the mean value of *D*_1_. Each point represents the mean value of *D*_1_ across 100 simulated datasets. b) Simulations show that variation in γ_2_ (color legend) changes the mean value of *D*_1_. Each point represents the mean value of *D*_1_ across 100 simulated datasets.

Finally, the results obtained from our two different simulation approaches are virtually identical (see Figure S1), confirming that they both reflect valid ways of simulating introgression.

### Power of *D*_2_ to determine the direction of introgression

We investigated the power of the *D*_2_ statistic to distinguish between different directions of introgression, and how this power is affected by the time between speciation events, *t*_2_ − *t*_1_, in addition to the two other parameters described above. Our model predicts, under our previously stated assumptions, that introgression in the C→B direction should produce values of *D*_2_ not different from 0, while introgression in the B→C direction should produce values significantly larger than 0. Furthermore, the magnitude of this difference should increase linearly as a function of *t*_2_ − *t*_1_, increasing the power of the test.

The results of our simulations, shown in Figure 6a, confirm the predictions of the model. In the B→C direction, the value of *D*_2_ increases linearly as a function of *t*_2_ − *t*_1_, whereas *D*_2_ values remain centered around 0 for introgression in the C→B direction. For the particular introgression scenario examined (introgression 0.6*N* generations after speciation), our statistic always has excellent power to distinguish between the two directions (Supplementary Tables S3 and S4). The trend of our simulated results suggests that as the time between speciation events continues to decrease, the power of the *D*_2_ statistic will be reduced. Therefore, our statistic may be prone to accepting the null hypothesis (C→B introgression) when two speciation events are followed very closely in time.

**Figure 6:**
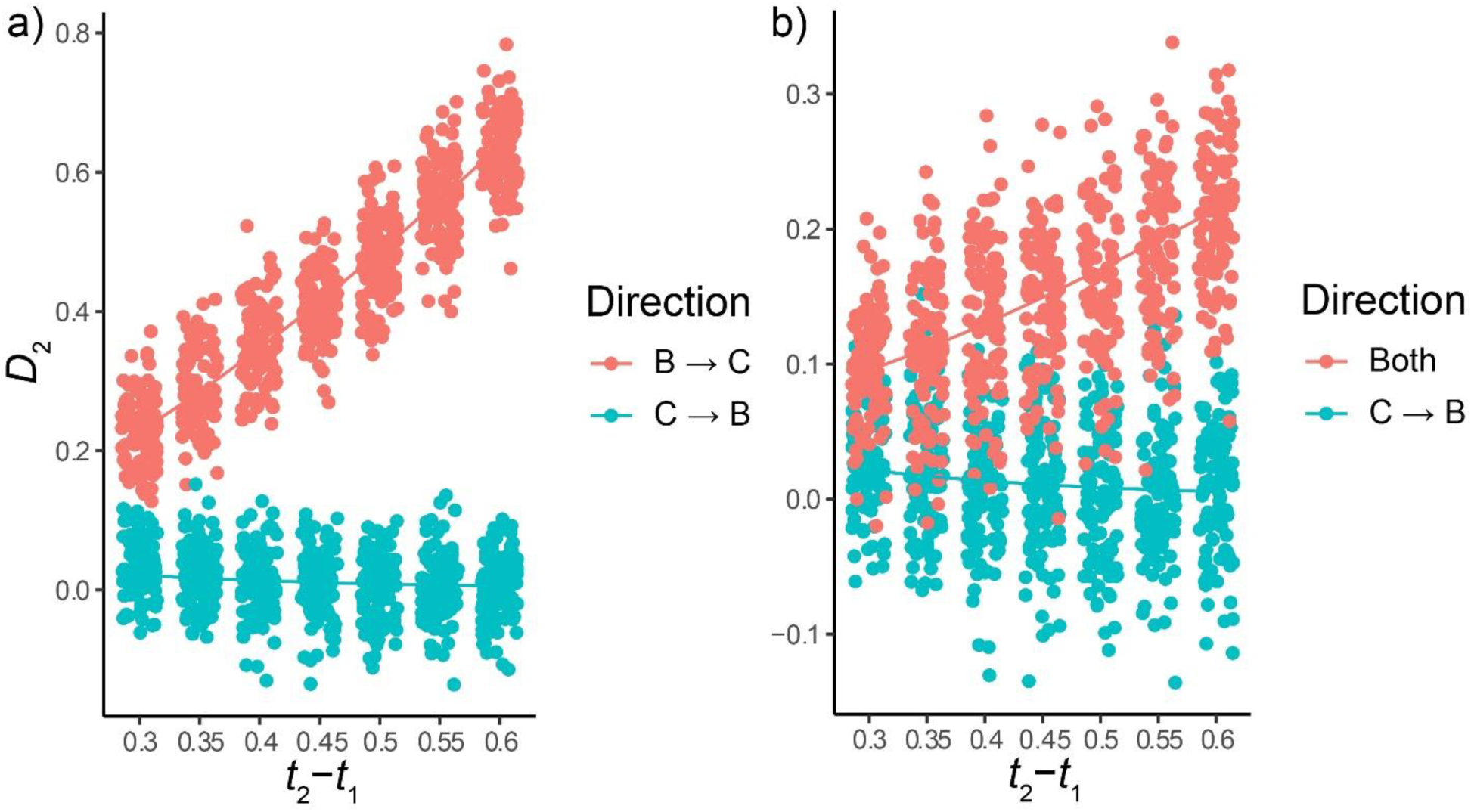
*D*_2_ as a function of the time between speciation events, *t*_2_ − *t*_1_. The color legend denotes the direction of introgression, with the solid line showing expected values from our model, and the points showing simulated values. Panel a) contrasts the two directions individually, while panel b) contrasts C→B introgression with introgression in both directions. Time on the x-axis is measured in units of 4*N*.

Introgression in both directions may reduce the power of *D*_2_ to reject the null, again similarly to the behavior of *D*_1_. This is due to the presence of ((B,C),A) gene trees concordant with parent tree 2, which share the same A-C divergence times as ((A,B),C) trees concordant with parent tree 1. We investigated this prediction, and how it interacts with the time between speciation events, with a set of simulations including equal contributions of parent trees 2 and 3, again 0.6*N* generations after speciation. The simulated results confirm the predictions of our model (Figure 6b). The power of the statistic to reject C→B introgression alone is reduced compared to the same values of *t*_2_ − *t*_1_ when introgression is B→C, but power may still be good if the time between speciation events is high enough (Figure 6b).

We also tested whether variation in *N*_1_ / *N*_2_ and γ_2_ would affect the *D*_2_ statistic in the same ways they affect *D*_1_. To do this, we simulated across the same range of parameters for the *D*_2_ statistic. The results of these simulations suggest that *D*_2_ is substantially more robust to variation in *N* than *D*_1_ (Figure 7a). Moderate false positive rates (incorrectly rejecting null of C → B introgression) of 10-20% begin to appear only when *N*_1_ is 2/3 of *N*_2_ or lower, and there appears to be no affect when *N*_1_ is larger than *N*_2_ (Supplementary Table S3). The statistic may be sensitive to false negatives (incorrectly accepting null of C → B) if *N*_1_ is lower than *N*_2_ and there is a short time between lineage-splitting events (ie. *t*_2_ − *t*_1_ is small); our simulations show a false negative rate of 34% when *N*_1_ is half of *N*_2_ and *t*_2_ − *t*_1_ is 0.5 (Supplementary Table S4). *D*_2_ also appears to be sensitive to variation in γ_2_ (or γ_3_, depending on the direction of introgression). Values of γ_2_ less than or equal to 0.2, or greater than or equal to 0.8, can lead to high false positive rates (Supplementary Table S3). Unlike *D*_1_, the false positive rate of *D*_2_ may be robust to some variation in γ_2_ if *t*_2_ − *t*_1_ is large; in our simulations, the false positive rate remains at or below standard significance at γ_2_ = 0.65 when *t*_2_ − *t*_1_ is 0.6 (Supplementary Table S3). *D*_2_ can also have a high false negative rate in certain regions of parameter space: this is again caused by the effects of γ_3_ negating the divergence time signal. Our simulations suggest that *D*_2_ is generally at a higher risk for false negatives when γ_3_ is small (0.35 or less), but this effect interacts with *t*_2_ − *t*_1_ in a somewhat complex way (Supplementary Table S4).

**Figure 7:**
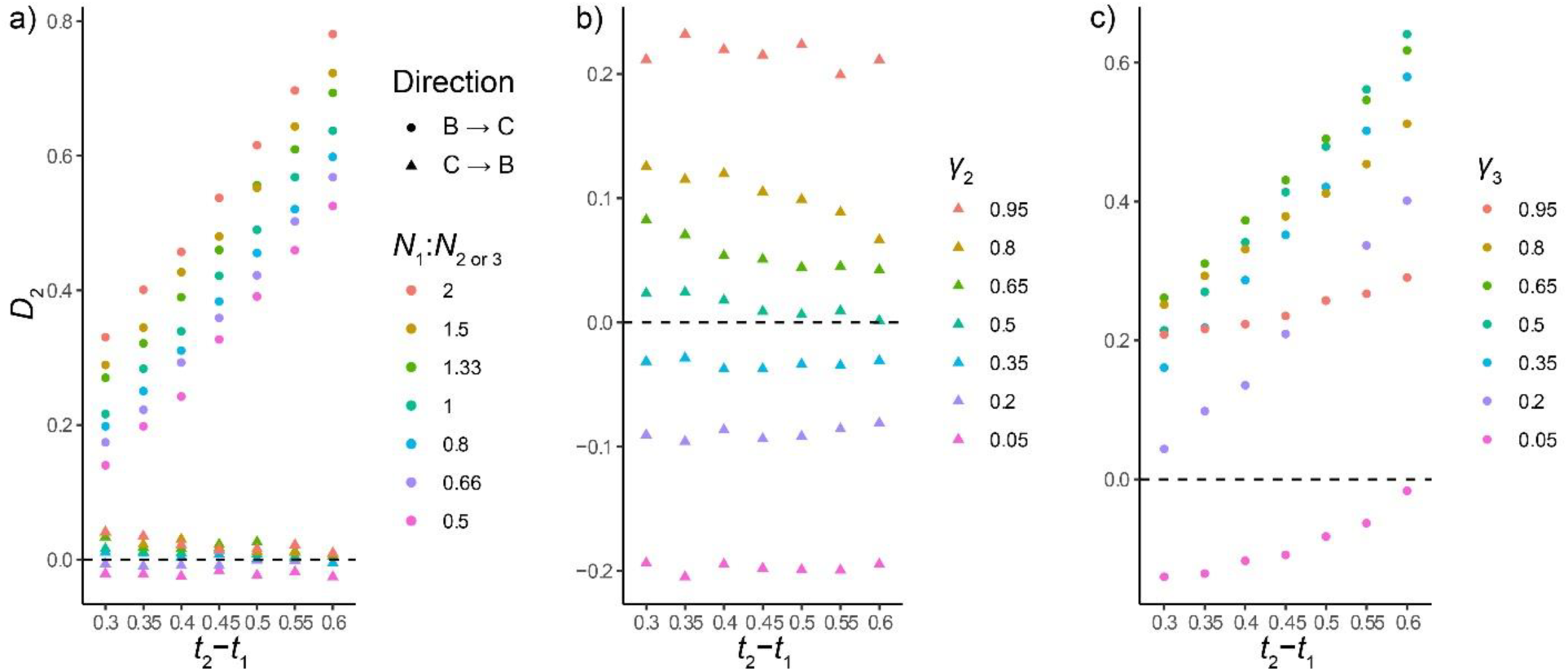
*D*_2_ as a function of variation in *N*_1_ / *N*_2_ and γ_2_. a) Effects of variation in *N* (color legend) on *D*_2_ for both directions of introgression (shape legend). Each point represents the mean of 100 simulated datasets. b) Effects of variation in γ_2_ (color legend) on *D*_2_ for C → B introgression. Each point represents the mean of 100 simulated datasets. c) Effects of variation in γ_3_ (color legend) on *D*_2_ for B → C introgression. Each point represents the mean of 100 simulated datasets.

Our simulation results highlight the fact that the signal the *D*_2_ statistic detects is that of B→C introgression, regardless of whether introgression also happened in the other direction. Therefore, a significantly positive value of *D*_2_ cannot explicitly distinguish between B→C introgression alone and B→C introgression coupled with some C→B introgression. Conversely, a non-significant value of *D*_2_ does not rule out the presence of some B→C introgression. As with *D*_1_, the relative magnitude of the contributions of parent trees 2 and 3 will affect this result, as will the timing between speciation events. Therefore, the most accurate way to interpret any value of *D*_2_ is to state the primary direction of introgression, rather than stating with certainty that introgression occurred only in one direction or another.

### Analysis of *S. paradoxus*

To demonstrate the use of our model to test the hypothesis of homoploid hybrid speciation, we calculated *D*_1_ from three lineages of the wild yeast *Saccharomyces paradoxus* and an outgroup and obtained a value of −0.0004. If the hypothesis that the *SpC*\* lineage of *S. paradoxus* is a hybrid species is correct (cf. Leducq et al. 2016), then we would expect *D*_1_ to follow the distribution obtained from our simulations under a homoploid hybrid speciation scenario. A *D*_1_ value significantly deviating from this expectation would indicate a bad fit to the HHS hypothesis.

To interpret our empirical estimate of *D*_1_ we simulated the *S. paradoxus* system under HHS. We used our estimated values of *θ* of 2.23 × 10−4 and 7.37 × 10−4 for the *SpC* and *SpB* populations, respectively, and values of the per-generation mutation rate and number of generations per year of 1.84 × 10−10 and 2920, respectively (Fay and Benavides 2005, Zhu et al. 2014). *N*_1_ and *N*_2_ were estimated at approximately 3 × 105 and 1 × 106, while divergence times in units of 4*N* generations were estimated at 112 and 20.2 for *t*_2_ and *t*_1_, respectively (see Supplementary Materials for details as to how all parameters were calculated.).

Using these population parameters, we simulated 10000 *D*_1_ statistics under a homoploid hybrid speciation scenario. Our simulations replicate the pattern observed in the data of a total absence of incomplete lineage sorting, with all trees coming from a single parent tree having a single topology. The simulations also show that under these parameters the value of *D*_1_ is expected be negative (Figure 8). However, the mean value of *D*_1_ = −0.0004 observed in the *S. paradoxus* system is highly unlikely to have arisen under our simulated hybrid speciation scenario (*P* < 1.0 × 10−4, rank significance test described in Methods; Figure 8). This result strongly rejects homoploid hybrid speciation for the *S. paradoxus* system.

**Figure 8:**
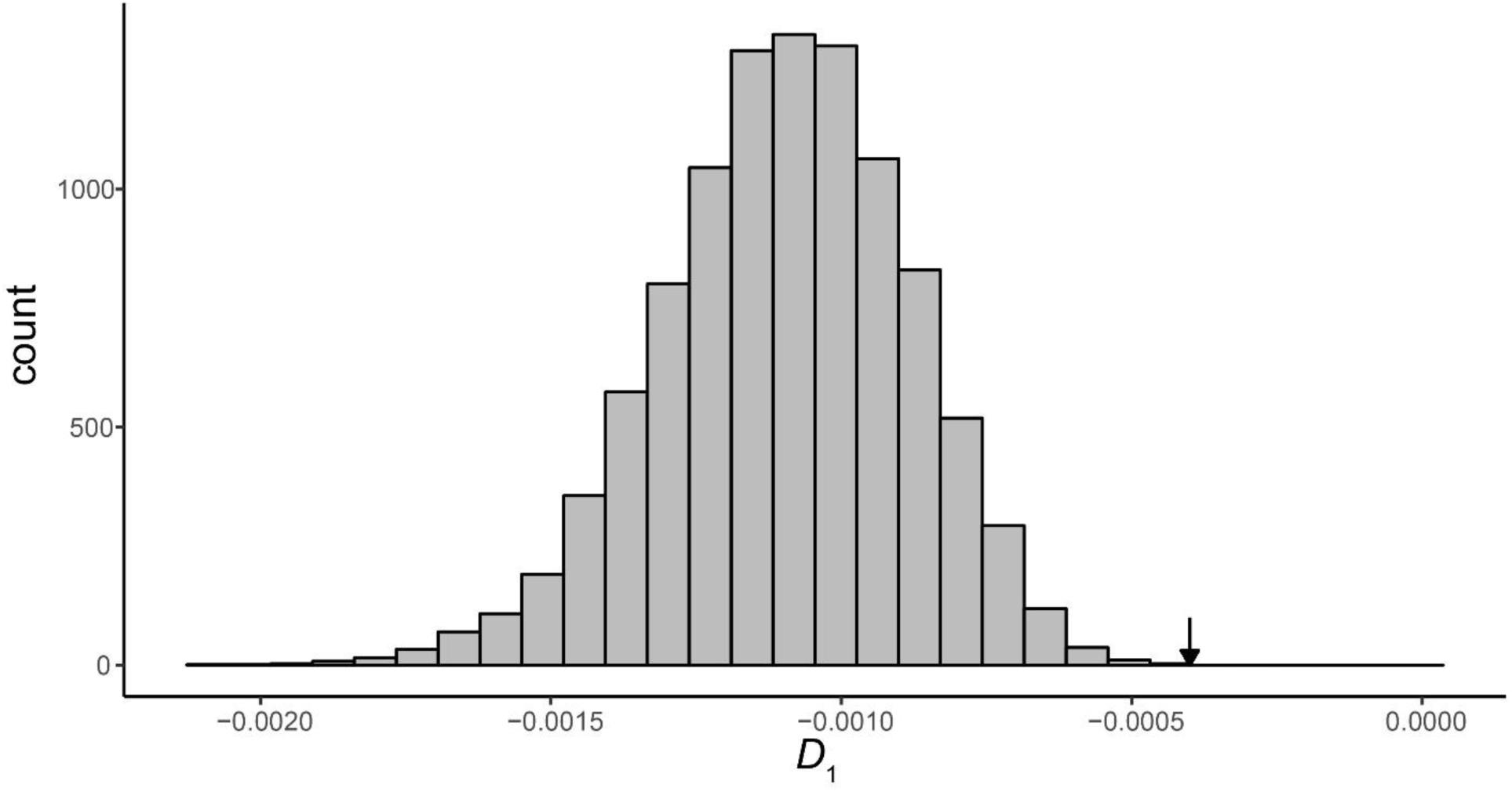
Distribution of 10,000 *D*_1_ statistics simulated under a hybrid speciation scenario using *S. paradoxus* demographic parameters. The arrow indicates the mean value of *D*_1_ estimated for the *S. paradoxus* system, which was significant by a rank significance test (*D*_1_ = −0.0004, p < 1×10−4).

## Discussion

There are now multiple methods that use only one sequence per lineage to detect the presence of gene flow between species (reviewed in Elworth et al. 2018). These methods all take advantage of the fact that expectations for the frequency of different gene tree topologies can easily be calculated under ILS, with deviations from these expectations often indicating the presence of gene flow. However, gene tree topologies alone, without branch lengths, cannot distinguish among various biological scenarios involving introgression (Zhu and Degnan 2017). In particular, the scenarios represented in Figure 1a and 1b cannot be distinguished solely using tree topologies, leading to general confusion and a proliferation of claims about “hybrid species.” The goal of the current study is to model possible histories of gene flow under the multispecies network coalescent, and to present two new test statistics to explicitly differentiate among such histories. While neither of these statistics should be used as a test for the presence of introgression itself, they complement other widely used statistics that can be used for this purpose and that depend on the same sampling scheme. In what follows we discuss the limitations and implications of this work.

### Limitations of our model

Our model of gene flow between lineages is simplified in multiple ways, and application to real data will often have to confront several assumptions we have made. We have assumed that a single, instantaneous introgression event leads to the generation of a single alternative history to the species tree. If instead there are multiple introgression events, each additional event will generate a new reticulation in the species network (and an additional parent tree), in the extreme producing infinitely many parent trees for continuous stretches of introgression. Under such scenarios we expect an increase in the variance in coalescent times, but still expect our statistics to capture the main history of gene flow (e.g. Figure 5b). One exception is discussed further in the next section when considering hypotheses about homoploid hybrid speciation.

We have modeled gene flow as a “horizontal” edge in the network, either because of post-speciation introgression (Figure 1a) or hybrid speciation (Figure 1b). In such representations the migrant individuals do not have an evolutionary history that is independent of the donor lineage. In contrast, the representation in Figure 1c uses “non-horizontal” edges to model gene flow, possibly indicating histories in which migrant individuals can evolve independently (Degnan 2018). Such histories could possibly indicate biological scenarios involving lineage fusion (e.g. Kearns et al. 2018) or where unsampled or extinct lineages are the donor population, but seem much less relevant to most gene flow events. However, this representation is commonly used in two settings. First, non-horizontal migration edges are often used to indicate the direction of introgression after speciation, with the first bifurcation in Figure 1c representing speciation between species A and B, and the second representing gene flow from C into B (e.g. Huson et al. 2010). This use does not imply an independent history, but instead simply the direction of introgression. Second, some methods for inferring the topology of a species network require or include non-horizontal edges (e.g. Yu et al. 2014; Solís-Lemus et al. 2017; Zhang et al. 2018). Although this choice adds parameters to these models, it also makes some calculations easier. Despite the computational convenience, this seems to generally be a less biologically realistic choice. Furthermore, the choice of horizontal edges, as is used here, makes a clear distinction between the species history and any introgression histories; models that use non-horizontal edges cannot distinguish between such alternatives (cf. Wen et al. 2016a).

Our model assumes that coalescence times and gene tree frequencies follow neutral expectations. In the presence of either hitchhiking or background selection, *N*_*e*_ will be reduced, reducing coalescence times and increasing the concordance of gene trees with their respective parent trees. This latter consequence should actually improve the power of our tests by reducing the incomplete lineage sorting that occurs within each parent tree. A further complication may be introduced because of the interaction between selection and introgression: in a number of systems introgression appears to occur more frequently in regions of the genome with higher recombination, which are less affected by linked selection (Geraldes et al. 2011; Brandvain et al. 2014; Aeschbacher et al. 2017; Schumer et al. 2018b). Because such regions will have larger *N_e_*, they may show more discordance and longer times to coalescence than average loci. In order to overcome such confounding factors, it may be best to make comparisons among genomic windows with similar recombination rates.

Finally, we have assumed that trees can be identified from individual non-recombining windows of the genome. In reality, such windows are likely to be very small (e.g. Mendes et al. 2018), and averaging multiple non-recombining windows together may lead to the reconstruction of an incorrect topology (Schierup and Hein 2000; Kubatko and Degnan 2007; Martin and Van Belleghem 2017). While this implies that genomic windows should not be too big, neighboring non-recombining segments will have correlated topologies, and inferred topologies from regions on the order of the length of single genes are not very different from the true topology (Lanier and Knowles 2012).

### Implications for homoploid hybrid speciation

It has become increasingly apparent that HHS is a process that can generate new diversity quickly, with several charismatic examples now known (e.g. in Darwin’s finches; Lamichhaney et al. 2018). Characterizing the frequency with which HHS occurs in nature is therefore important for our understanding of speciation and the evolution of reproductive isolation, though strict criteria for identifying true cases of HHS have been lacking. Schumer et al. (2014) proposed three pieces of evidence that are required to demonstrate HHS: 1) evidence of introgression, 2) evidence of reproductive isolation of the hybrid lineage from both parents, 3) evidence of a causal link between introgression and reproductive isolation. While relatively standard methods exist for evaluating criteria 1 and 2, it is much more difficult to explicitly evaluate criterion 3. Our *D*_1_ statistic is unique in that it has a specific distribution of expected values under a hybrid speciation scenario, which can be predicted precisely using modelling and/or simulation. Therefore, it provides an explicit test of criterion 3 by asking whether speciation and introgression are effectively simultaneous. Such a relationship would strongly imply a causal link.

A commonly employed expectation for HHS is that there should be an approximately 50:50 split of two contrasting histories in the genome of the hybrid, as would be expected if each parent species contributed equally. However, this pattern may be misleading for at least two reasons. First, not all hybrid species are the result of isolation caused in the F1 generation of crosses between two species. For example, the hybrid butterfly *Heliconius heurippa* likely arose through two generations of backcrossing, resulting in an 82.5:12.5 pattern of ancestry (Mavarez et al. 2006). Selection or drift may also cause deviations from 50:50 expectations in cases of true HHS. Second, introgression without hybrid speciation can be extensive, affecting 50% of the genome or more (e.g. in *Anopheles* mosquitoes; Fontaine et al. 2015; Wen et al. 2016a). Our *D*_1_ statistic overcomes this limitation by explicitly allowing the admixture proportion to vary when predicting its expected value under an HHS scenario.

There are several other biological scenarios in which the *D*_1_ statistic in particular may be misleading, resulting in either incorrect rejection or acceptance of HHS. If hybrid speciation is followed by extinction of one parent lineage, then one sampled taxon will be more distantly related to the hybrid than the other; this will lead to values of *D*_1_ inconsistent with HHS even though it has occurred. Similarly, if introgression occurs after hybrid speciation, the value of *D*_1_ could be dominated by the more recent event, again leading to false rejection of HHS. While both of these scenarios are problems for *D*_1_, they would also be problems for any other methods attempting to distinguish HHS from introgression-after-speciation. Lastly, if introgression has occurred shortly after speciation, but is not causally related to it, there simply may not be enough signal in the data to distinguish this scenario from one in which they are simultaneous.

### Application of D_1_ to an empirical example in yeast

One clade of the yeast species *Saccharomyces paradoxus* (denoted *SpC*\*) has been proposed to be a homoploid hybrid (Leducq et al. 2016). Our estimate of *D*_1_ from genomic data suggests that it is highly unlikely to have arisen under a hybrid speciation scenario; here, we discuss the implications of this result for the system and for attempts to infer HHS in general.

In their analysis of the genome of *SpC\**, Leducq et al. (2016) concluded that each locus could be classified into one of two topologies: either [((*SpC*\*,*SpC*),*SpB*)], which comprises 92–97% of the genome, and [((*SpC*\*,*SpB*),*SpC*)], which comprises the remainder. The absence of a third gene tree topology, [((*SpC*,*SpB*),*SpC\**)], leads us to believe that incomplete lineage sorting is minimal or entirely absent in this system. Therefore, this represents a unique case in which the genome is comprised of two specific gene trees whose estimated coalescence times can be compared directly: a gene tree concordant with the species history (as described by equation 1) and a gene tree concordant with a history of introgression (as described by equation 9). Our simulations using population parameters from the *S. paradoxus* system replicate the observation that incomplete lineage sorting is absent from the data.

Given the results of our analysis, the simplest explanation for the observed pattern is that there was introgression between *SpC*\* and *SpB* after the split of *SpC* and *SpC\**. It is also possible that *SpC*\* originated as a hybrid species that has since undergone further introgression only with *SpB*. While it may be difficult to distinguish these two hypotheses, it is clear from our results that the primary signal carried in the data is one of introgression following speciation, rather than homoploid hybrid speciation.

### Considerations for the D_2_ statistic

We have also introduced the *D*_2_ statistic for distinguishing the direction of introgression between two taxa (see Rouard et al. 2018 for an empirical example using this test). Despite the importance of understanding the direction of gene flow, relatively few studies have explicitly attempted to provide a solution to the problem when only a single sequence is sampled from each lineage. Pease & Hahn (2015) developed a set of *D*_FOIL_ statistics which can infer the direction of introgression in such cases; however, these statistics require information from four ingroup taxa, and the taxa in question must have a symmetrical tree topology. These considerations limit the generality of *D*_FOIL_ statistics.

The *D*_2_ statistic can be calculated from three ingroup taxa and an outgroup; this is similar to the sampling used for the original *D* statistic (Green et al. 2010) and other related tests using gene tree topologies (e.g. Huson et al. 2005). Using only the frequencies of topologies (or nucleotide site patterns that reflect these underlying topologies), such tests cannot distinguish the direction of introgression. While the internal branches on the two major tree topologies produced by alternative directions of introgression do differ in length (i.e. from *t*_2_ to *t*_m_ in parent tree 2 vs. from *t*_1_ to *t*_m_ in parent tree 3; Figure 2), this difference is not detectable using the *D* statistic (Martin et al. 2015). There are of course population genetic methods for inferring the direction of gene flow, and these can even be used between pairs of sister species (e.g. Lohse et al. 2016). However, these methods both require more sequences from each taxon and are more sensitive to violations of their assumptions due to linked selection because they are more dependent on the variance in coalescence times. *D*_2_ also appears to be more robust than *D*_1_ to variation in *N* between populations (Figure 7a). Although such variation can be accounted for in simulations, this robustness makes it less important that accurate estimates of *N* be made.

### Conclusions

Here, we have developed a network coalescent model that can predict coalescence times generated under various introgression scenarios. From this model, we propose two new test statistics, named *D*_1_ and *D*_2_, which can be used to test hypotheses about the relative timing and direction of introgression. *D*_1_ evaluates the null hypothesis that lineage-splitting and introgression occur simultaneously, which is expected to occur during homoploid hybrid speciation or in the creation of admixed populations. This statistic builds on descriptive models by providing a quantitative means for addressing hypotheses related to HHS. *D*_2_ is designed to be a test of the null hypothesis that introgression occurred primarily in the C→B direction (from an unpaired species to a paired species), with rejection of the null indicating that introgression primarily occurred in the B→C direction. Our model and statistics can be used with simulated data to provide powerful hypothesis testing in a variety of systems; this is highlighted in our application of the *D*_1_ statistic to an empirical dataset from wild yeast.

## Supporting information

## Acknowledgements

We thank Nick Barton and two referees for their very helpful comments, as well as Christian Landry and Jean-Baptiste Leducq for their assistance and for sharing their data with us.

Discussions with Rafael Guerrero, Fabio Mendes, and Ben Rosenzweig also helped to improve the work. This research was supported by National Science Foundation grant MCB-1127059 to M.W.H.

