## Supplementary material for "The timing and direction of introgression under the multispecies network coalescent"

Supplementary Materials and Methods

Mark S. Hibbins* and Matthew W. Hahn*†

*Department of Biology and †Department of Computer Science
Indiana University, Bloomington IN 47405

November 20, 2018

#### Calculation of expectations for $D_1$

The $D_1$ statistic is defined, under the assumption that only parent trees 1 and 2 are relevant, in terms of coalescent expectations as:

$$E[D_1] = (f_{AB1_1}E[t_{A-B}|AB1_1] + f_{AB2_1}E[t_{A-B}|AB2_1] + f_{AB_2}E[t_{A-B}|AB_2]) - (f_{BC1_2}E[t_{B-C}|BC1_2] + f_{BC2_2}E[t_{B-C}|BC2_2] + f_{BC_1}E[t_{B-C}|BC_1]) \quad (1)$$

(This is equation 18 in the main text). If we consider coalescent time in units of $2N$ generations, substituting each time term with its mathematical expectation yields the following:

$$E[D_1] = (f_{AB1_1}(t_1 + (1 - \frac{t_2 - t_1}{e^{t_2 - t_1} - 1})) + f_{AB2_1}(t_2 + \frac{1}{3}) + f_{AB_2}(t_2 + \frac{1}{3}) - (f_{BC1_2}(t_{\mathrm{m}} + (1 - \frac{t_2 - t_{\mathrm{m}}}{e^{t_2 - t_{\mathrm{m}}} - 1})) + f_{BC2_2}(t_2 + \frac{1}{3}) + f_{BC_1}(t_2 + \frac{1}{3})) \quad (2)$$

It is important to note that, due to the conditional nature with which our statistic is defined, the frequency terms in equations S.1 and S.2 cannot simply be taken as gene tree frequency expectations from individual parent trees. Rather, each component is conditional on particular gene tree topologies, which can come from different parent trees. Therefore, we must: 1) weight each frequency by the fraction of loci in the genome that evolved within that parent history; 2) normalize each frequency so that it is with respect to all the gene trees of a particular topology, rather than with respect to all gene trees in the dataset.

First, the expected frequencies of each genealogy within a parent tree are a classical result from the coalescent:

$$P(\text{concordance}|\text{lineage sorting}) = 1 - e^{-\tau} \quad (3)$$

$$P(\text{concordance}|\text{ILS}) = P(\text{discordance}|\text{ILS}) = \frac{1}{3}e^{-\tau} \quad (4)$$

Where $\tau$ represents the internal branch lengths — that is, the time between lineage splitting events (i.e. between $t_1$, $t_2$, and $t_{\mathrm{m}}$) — in each parent tree. Equation S.3 represents the expected frequency for gene trees $AB1_1$, $BC1_2$, and $BC1_3$ in our model, while equation S.4 can be used as the frequency for the other gene trees (all conditional on their parent trees).

Let us define the percentage contribution of parent tree 2 to the history of the species we are considering as $\gamma$, and the contribution from parent tree 1 as $1-\gamma$. To normalize each gene tree frequency with respect to all the trees sharing its topology, we multiply the expected frequency given above by the parental contribution of that gene tree, divided by the sum of the expectation for all frequencies sharing its topology. With these two considerations, and substituting the appropriate times from our model for $\tau$, we can define each of the frequency terms used in our statistic:

$$f_{AB1_1} = \frac{(1-\gamma)(1-e^{-(t_2-t_1)})}{(1-\gamma)(1-e^{-(t_2-t_1)})+(1-\gamma)\frac{1}{3}e^{-(t_2-t_1)}+\gamma\frac{1}{3}e^{-(t_2-t_\text{m})}} \tag{5}$$

$$f_{AB2_1} = \frac{(1-\gamma)\frac{1}{3}e^{-(t_2-t_1)}}{(1-\gamma)(1-e^{-(t_2-t_1)})+(1-\gamma)\frac{1}{3}e^{-(t_2-t_1)}+\gamma\frac{1}{3}e^{-(t_2-t_\text{m})}} \tag{6}$$

$$f_{AB_2} = \frac{\gamma\frac{1}{3}e^{-(t_2-t_\text{m})}}{(1-\gamma)(1-e^{-(t_2-t_1)})+(1-\gamma)\frac{1}{3}e^{-(t_2-t_1)}+\gamma\frac{1}{3}e^{-(t_2-t_\text{m})}} \tag{7}$$

$$f_{BC1_2} = \frac{\gamma(1-e^{-(t_2-t_\text{m})})}{\gamma(1-e^{-(t_2-t_\text{m})})+\gamma\frac{1}{3}e^{-(t_2-t_\text{m})}+(1-\gamma)\frac{1}{3}e^{-(t_2-t_1)}} \tag{8}$$

$$f_{BC2_2} = \frac{\gamma\frac{1}{3}e^{-(t_2-t_\text{m})}}{\gamma(1-e^{-(t_2-t_\text{m})})+\gamma\frac{1}{3}e^{-(t_2-t_\text{m})}+(1-\gamma)\frac{1}{3}e^{-(t_2-t_1)}} \tag{9}$$

$$f_{BC_1} = \frac{(1-\gamma)\frac{1}{3}e^{-(t_2-t_1)}}{\gamma(1-e^{-(t_2-t_\text{m})})+\gamma\frac{1}{3}e^{-(t_2-t_\text{m})}+(1-\gamma)\frac{1}{3}e^{-(t_2-t_1)}} \tag{10}$$

In the special case where $t_1-t_\text{m}=0$, all terms in equation S.5 exactly cancel out, leaving us with $E[D_1]=0$, as stated in the main text. The expected value of $D_1$ is a linear function of $t_1-t_\text{m}$.

We also considered values of $D_1$ under a scenario in which introgression has occurred in both directions. The expectation of $D_1$ in this circumstance must also now include the contributions from parent tree 3 in addition to 2. With this additional consideration of the gene tree frequencies from parent tree 3, the expectation for $D_1$ is as follows:

$$\begin{aligned}E[D_1] = (f_{AB1_1}(t_1+(1-\frac{t_2-t_1}{e^{t_2-t_1}-1}))+f_{AB2_1}(t_2+\frac{1}{3})+f_{AB_2}(t_2+\frac{1}{3})+f_{AB_3}(t_1+\frac{1}{3})\\ -(f_{BC1_2}(t_\text{m}+(1-\frac{t_2-t_\text{m}}{e^{t_2-t_\text{m}}-1}))+f_{BC2_2}(t_2+\frac{1}{3})+f_{BC_1}(t_2+\frac{1}{3})\\ +f_{BC1_3}(t_\text{m}+(1-\frac{t_1-t_\text{m}}{e^{t_1-t_\text{m}}}))+f_{BC2_3}(t_1+\frac{1}{3}))\end{aligned} \tag{11}$$

Our normalized gene tree frequencies must now also account for the presence of these additional gene trees. We define $\gamma_2$ as the contribution of parent tree 2, $\gamma_3$ as the contribution of parent tree 3, and $1-\gamma_2-\gamma_3$ as the contribution of parent tree 1. Using the same normalization approach as above, we obtain:

$$f_{AB1_1} = \frac{(1-\gamma_2-\gamma_3)(1-e^{-(t_2-t_1)})}{(1-\gamma_2-\gamma_3)(1-e^{-(t_2-t_1)})+(1-\gamma_2-\gamma_3)\frac{1}{3}e^{-(t_2-t_1)}+\gamma_2\frac{1}{3}e^{-(t_2-t_\text{m})}+\gamma_3\frac{1}{3}e^{-(t_1-t_\text{m})}} \tag{12}$$

$$f_{AB2_1} = \frac{(1-\gamma_2-\gamma_3)\frac{1}{3}e^{-(t_2-t_1)}}{(1-\gamma_2-\gamma_3)(1-e^{-(t_2-t_1)})+(1-\gamma_2-\gamma_3)\frac{1}{3}e^{-(t_2-t_1)}+\gamma_2\frac{1}{3}e^{-(t_2-t_\text{m})}+\gamma_3\frac{1}{3}e^{-(t_1-t_\text{m})}} \tag{13}$$

$$f_{AB_2} = \frac{\gamma_2\frac{1}{3}e^{-(t_2-t_\text{m})}}{(1-\gamma_2-\gamma_3)(1-e^{-(t_2-t_1)})+(1-\gamma_2-\gamma_3)\frac{1}{3}e^{-(t_2-t_1)}+\gamma_2\frac{1}{3}e^{-(t_2-t_\text{m})}+\gamma_3\frac{1}{3}e^{-(t_1-t_\text{m})}} \tag{14}$$

$$f_{AB_3} = \frac{\gamma_3 \frac{1}{3} e^{-(t_1 - t_{\mathrm{m}})}}{(1-\gamma_2-\gamma_3)(1-e^{-(t_2-t_1)}) + (1-\gamma_2-\gamma_3)\frac{1}{3}e^{-(t_2-t_1)} + \gamma_2 \frac{1}{3} e^{-(t_2-t_{\mathrm{m}})} + \gamma_3 \frac{1}{3} e^{-(t_1-t_{\mathrm{m}})}} \tag{15}$$

$$f_{BC1_2} = \frac{\gamma_2(1-e^{-(t_2-t_{\mathrm{m}})})}{\gamma_2(1-e^{-(t_2-t_{\mathrm{m}})}) + \gamma_2(\frac{1}{3}e^{-(t_2-t_{\mathrm{m}})}) + (1-\gamma_2-\gamma_3)(\frac{1}{3}e^{-(t_2-t_1)}) + \gamma_3(1-e^{-(t_1-t_{\mathrm{m}})}) + \gamma_3(\frac{1}{3}e^{-(t_1-t_{\mathrm{m}})})} \tag{16}$$

$$f_{BC2_2} = \frac{\gamma_2(\frac{1}{3}e^{-(t_2-t_{\mathrm{m}})})}{\gamma_2(1-e^{-(t_2-t_{\mathrm{m}})}) + \gamma_2(\frac{1}{3}e^{-(t_2-t_{\mathrm{m}})}) + (1-\gamma_2-\gamma_3)(\frac{1}{3}e^{-(t_2-t_1)}) + \gamma_3(1-e^{-(t_1-t_{\mathrm{m}})}) + \gamma_3(\frac{1}{3}e^{-(t_1-t_{\mathrm{m}})})} \tag{17}$$

$$f_{BC_1} = \frac{(1-\gamma_2-\gamma_3)(\frac{1}{3}e^{-(t_2-t_1)})}{\gamma_2(1-e^{-(t_2-t_{\mathrm{m}})}) + \gamma_2(\frac{1}{3}e^{-(t_2-t_{\mathrm{m}})}) + (1-\gamma_2-\gamma_3)(\frac{1}{3}e^{-(t_2-t_1)}) + \gamma_3(1-e^{-(t_1-t_{\mathrm{m}})}) + \gamma_3(\frac{1}{3}e^{-(t_1-t_{\mathrm{m}})})} \tag{18}$$

$$f_{BC1_3} = \frac{\gamma_3(1-e^{-(t_1-t_{\mathrm{m}})})}{\gamma_2(1-e^{-(t_2-t_{\mathrm{m}})}) + \gamma_2(\frac{1}{3}e^{-(t_2-t_{\mathrm{m}})}) + (1-\gamma_2-\gamma_3)(\frac{1}{3}e^{-(t_2-t_1)}) + \gamma_3(1-e^{-(t_1-t_{\mathrm{m}})}) + \gamma_3(\frac{1}{3}e^{-(t_1-t_{\mathrm{m}})})} \tag{19}$$

$$f_{BC2_3} = \frac{\gamma_3(\frac{1}{3}e^{-(t_1-t_{\mathrm{m}})})}{\gamma_2(1-e^{-(t_2-t_{\mathrm{m}})}) + \gamma_2(\frac{1}{3}e^{-(t_2-t_{\mathrm{m}})}) + (1-\gamma_2-\gamma_3)(\frac{1}{3}e^{-(t_2-t_1)}) + \gamma_3(1-e^{-(t_1-t_{\mathrm{m}})}) + \gamma_3(\frac{1}{3}e^{-(t_1-t_{\mathrm{m}})})} \tag{20}$$

### Calculation of expectations for $D_2$

Using the same conventions described above for $D_1$, we can define the expectation of $D_2$ in each introgression scenario. For the direction $C \to B$:

$$\begin{aligned} E[D_2|C \to B] = (f_{AB1_1}E[t_{A-C}|AB1_1] + f_{AB2_1}E[t_{A-C}|AB2_1] + f_{AB_2}E[t_{A-C}|AB_2]) \\ - (f_{BC1_2}E[t_{A-C}|BC1_2] + f_{BC2_2}E[t_{A-C}|BC2_2] + f_{BC_1}E[t_{A-C}|BC_1]) \end{aligned} \tag{21}$$

This corresponds to equation 20 in the main text. Substituting the expected coalescent times:

$$\begin{aligned} E[D_2|C \to B] = (f_{AB1_1}(t_2+1)] + f_{AB2_1}(t_2+\frac{1}{3}+1) + f_{AB_2}(t_2+\frac{1}{3}+1) \\ - (f_{BC1_2}(t_2+1) + f_{BC2_2}(t_2+\frac{1}{3}+1) + f_{BC_1}(t_2+\frac{1}{3}+1)) \end{aligned} \tag{22}$$

The normalized gene tree frequencies used in S.21 and S.22 are defined above in equations S.5 - S.10. For the direction $B \to C$ we have:

$$\begin{aligned} E[D_2|B \to C] = (f_{AB1_1}E[t_{A-C}|AB1_1] + f_{AB2_1}E[t_{A-C}|AB2_1] + f_{AB_3}E[t_{A-C}|AB_3]) \\ - (f_{BC1_3}E[t_{A-C}|BC1_3] + f_{BC2_3}E[t_{A-C}|BC2_3] + f_{BC_1}E[t_{A-C}|BC_1]) \end{aligned} \tag{23}$$

Corresponding to equation 21 in the main text. Substituting:

$$\begin{aligned} E[D_2|B \to C] = (f_{AB1_1}(t_2+1)] + f_{AB2_1}(t_2+\frac{1}{3}+1) + f_{AB_3}(t_1+\frac{1}{3}+1) \\ - (f_{BC1_3}(t_1+1) + f_{BC2_3}(t_1+\frac{1}{3}+1) + f_{BC_1}(t_2+\frac{1}{3}+1)) \end{aligned} \tag{24}$$

Here, we need to define new normalized gene tree frequencies for the case when only gene trees from parent trees 1 and 3 are present. They are as follows:

$$f_{AB1_1} = \frac{(1-\gamma)(1-e^{-(t_2-t_1)})}{(1-\gamma)(1-e^{-(t_2-t_1)})+(1-\gamma)(\frac{1}{3}e^{-(t_2-t_1)})+\gamma(\frac{1}{3}e^{-(t_1-t_{\mathrm{m}})})} \tag{25}$$

$$f_{AB2_1} = \frac{(1-\gamma)(\frac{1}{3}e^{-(t_2-t_1)})}{(1-\gamma)(1-e^{-(t_2-t_1)})+(1-\gamma)(\frac{1}{3}e^{-(t_2-t_1)})+\gamma(\frac{1}{3}e^{-(t_1-t_{\mathrm{m}})})} \tag{26}$$

$$f_{AB_3} = \frac{\gamma(\frac{1}{3}e^{-(t_1-t_{\mathrm{m}})})}{(1-\gamma)(1-e^{-(t_2-t_1)})+(1-\gamma)(\frac{1}{3}e^{-(t_2-t_1)})+\gamma(\frac{1}{3}e^{-(t_1-t_{\mathrm{m}})})} \tag{27}$$

$$f_{BC1_3} = \frac{\gamma(1-e^{-(t_1-t_{\mathrm{m}})})}{\gamma(1-e^{-(t_1-t_{\mathrm{m}})})+\gamma(\frac{1}{3}e^{-(t_1-t_{\mathrm{m}})})+(1-\gamma)\frac{1}{3}e^{-(t_2-t_1)}} \tag{28}$$

$$f_{BC2_3} = \frac{\gamma(\frac{1}{3}e^{-(t_1-t_{\mathrm{m}})})}{\gamma(1-e^{-(t_1-t_{\mathrm{m}})})+\gamma(\frac{1}{3}e^{-(t_1-t_{\mathrm{m}})})+(1-\gamma)\frac{1}{3}e^{-(t_2-t_1)}} \tag{29}$$

$$f_{BC_1} = \frac{(1-\gamma)\frac{1}{3}e^{-(t_2-t_1)}}{\gamma(1-e^{-(t_1-t_{\mathrm{m}})})+\gamma(\frac{1}{3}e^{-(t_1-t_{\mathrm{m}})})+(1-\gamma)\frac{1}{3}e^{-(t_2-t_1)}} \tag{30}$$

Lastly, we define the expectation of $D_2$ when introgression occurs in both directions:

$$\begin{aligned} E[D_2|C\to B, B\to C] = {} & (f_{AB1_1}(t_2+1)+f_{AB2_1}(t_2+\frac{1}{3}+1)+f_{AB_2}(t_2+\frac{1}{3}+1)+f_{AB_3}(t_1+\frac{1}{3}+1)) \\ & -(f_{BC1_2}(t_2+1)+f_{BC2_2}(t_2+\frac{1}{3}+1)+f_{BC_1}(t_2+\frac{1}{3}+1)+f_{BC1_3}(t_1+1)+f_{BC2_3}(t_1+\frac{1}{3}+1)) \end{aligned} \tag{31}$$

Equation S.31 uses the gene tree frequencies defined in S.12 - S.20.

### $D_1$ Simulations

To begin, we performed a small set of simulations across 7 different combinations of parameters. This was for 3 reasons; 1) to check the simulated values against the expectations from our model; 2) to see how introgression in both directions affects the values of the statistics, and 3) to verify that our two methods of simulating introgression agree with one another. For the method where parent trees were simulated separately and then combined, we used the following commands:

For parent tree 1:

ms 4 1000 -T -I 4 1 1 1 1 -ej 4.0 2 1 -ej 0.6 3 2 -ej 0.3 4 3

For parent tree 2:

ms 4 1000 -T -I 4 1 1 1 1 -ej 4.0 2 1 -ej 0.6 4 2 -ej 0.3 3 2
ms 4 1000 -T -I 4 1 1 1 1 -ej 4.0 2 1 -ej 0.6 4 2 -ej 0.25 3 2
ms 4 1000 -T -I 4 1 1 1 1 -ej 4.0 2 1 -ej 0.6 4 2 -ej 0.2 3 2
ms 4 1000 -T -I 4 1 1 1 1 -ej 4.0 2 1 -ej 0.6 4 2 -ej 0.15 3 2
ms 4 1000 -T -I 4 1 1 1 1 -ej 4.0 2 1 -ej 0.6 4 2 -ej 0.1 3 2
ms 4 1000 -T -I 4 1 1 1 1 -ej 4.0 2 1 -ej 0.6 4 2 -ej 0.05 3 2
ms 4 1000 -T -I 4 1 1 1 1 -ej 4.0 2 1 -ej 0.6 4 2 -ej 0.0001 3 2

For parent tree 3:

ms 4 1000 -T -I 4 1 1 1 1 -ej 4.0 2 1 -ej 0.3 4 2 -ej 0.2999 3 2
ms 4 1000 -T -I 4 1 1 1 1 -ej 4.0 2 1 -ej 0.3 4 2 -ej 0.25 3 2
ms 4 1000 -T -I 4 1 1 1 1 -ej 4.0 2 1 -ej 0.3 4 2 -ej 0.2 3 2
ms 4 1000 -T -I 4 1 1 1 1- ej 4.0 2 1 -ej 0.3 4 2 -ej 0.15 3 2
ms 4 1000 -T -I 4 1 1 1 1 -ej 4.0 2 1 -ej 0.3 4 2 -ej 0.1 3 2
ms 4 1000 -T -I 4 1 1 1 1 -ej 4.0 2 1 -ej 0.3 4 2 -ej 0.05 3 2
ms 4 1000 -T -I 4 1 1 1 1 -ej 4.0 2 1 -ej 0.3 4 2 -ej 0.0001 3 2

For introgression in the $C \rightarrow B$ direction only, we used $\gamma_2 = 0.5$. For introgression in both directions, we used $\gamma_2 = \gamma_3 = 0.25$. We simulated introgression using the population split/rejoin method for the $C \rightarrow B$ direction only. This was done using the commands:

ms 4 2000 -T -I 4 1 1 1 1 -ej 4.0 2 1 -ej 0.6 3 2 -ej 0.3 4 3 -es 0.2999 3 0.5 -ej 0.2999 5 2
ms 4 2000 -T -I 4 1 1 1 1 -ej 4.0 2 1 -ej 0.6 3 2 -ej 0.3 4 3 -es 0.25 3 0.5 -ej 0.25 5 2
ms 4 2000 -T -I 4 1 1 1 1 -ej 4.0 2 1 -ej 0.6 3 2 -ej 0.3 4 3 -es 0.2 3 0.5 -ej 0.2 5 2
ms 4 2000 -T -I 4 1 1 1 1 -ej 4.0 2 1 -ej 0.6 3 2 -ej 0.3 4 3 -es 0.15 3 0.5 -ej 0.15 5 2
ms 4 2000 -T -I 4 1 1 1 1 -ej 4.0 2 1 -ej 0.6 3 2 -ej 0.3 4 3 -es 0.1 3 0.5 -ej 0.1 5 2
ms 4 2000 -T -I 4 1 1 1 1 -ej 4.0 2 1 -ej 0.6 3 2 -ej 0.3 4 3 -es 0.05 3 0.5 -ej 0.05 5 2
ms 4 2000 -T -I 4 1 1 1 1 -ej 4.0 2 1 -ej 0.6 3 2 -ej 0.3 4 3 -es 0.0001 3 0.5 -ej 0.0001 5 2

#### Simulating values of $N_2$

To simulate different values of $N_2$, we specified a population expansion / bottleneck within the internal branch of parent tree 2. Parent trees 1 and 2 were simulated separately and then combined into a single file for downstream analysis, with an admixture proportion $\gamma_2 = 0.5$. We simulated the $C \rightarrow B$ direction only. We simulated parent tree 1 using the command:

ms 4 49000 -t 0.01 -T -I 4 1 1 1 1 -ej 4.0 2 1 -ej 0.6 3 2 -ej 0.2999 4 3

And parent tree 2 using the command:

ms 4 49000 -t 0.01 -T -I 4 1 1 1 1 -ej 4.0 2 1 -ej 0.6 4 2 -ej tbs 3 2 -en tbs 2 tbs -en 0.6 2 1

And passed parent tree 2 the following combinations of parameters:

0.2999 0.2999 2
0.2999 0.2999 1.5
0.2999 0.2999 1.25
0.2999 0.2999 1
0.2999 0.2999 0.75
0.2999 0.2999 0.66
0.2999 0.2999 0.5
0.25 0.25 2
0.25 0.25 1.5
0.25 0.25 1.25
0.25 0.25 1
0.25 0.25 0.75
0.25 0.25 0.66
0.25 0.25 0.5

0.2 0.2 2
0.2 0.2 1.5
0.2 0.2 1.25
0.2 0.2 1
0.2 0.2 0.75
0.2 0.2 0.66
0.2 0.2 0.5
0.15 0.15 2
0.15 0.15 1.5
0.15 0.15 1.25
0.15 0.15 1
0.15 0.15 0.75
0.15 0.15 0.66
0.15 0.15 0.5
0.1 0.1 2
0.1 0.1 1.5
0.1 0.1 1.25
0.1 0.1 1
0.1 0.1 0.75
0.1 0.1 0.66
0.1 0.1 0.5
0.05 0.05 2
0.05 0.05 1.5
0.05 0.05 1.25
0.05 0.05 1
0.05 0.05 0.75
0.05 0.05 0.66
0.05 0.05 0.5
0.0001 0.0001 2
0.0001 0.0001 1.5
0.0001 0.0001 1.25
0.0001 0.0001 1
0.0001 0.0001 0.75
0.0001 0.0001 0.66
0.0001 0.0001 0.5

#### *Simulating values of* $\gamma_2$

We simulated across different values of $\gamma_2$ using the population split/join approach, as it allows the ancestry proportion to be varied directly. We simulated the $C \rightarrow B$ direction of introgression only, with a constant value of $N$ across the tree. We used the following command:

ms 4 98000 -t 0.01 -T -I 4 1 1 1 1 -ej 4.0 2 1 -ej 0.6 3 2 -ej 0.3 4 3 -es tbs 3 tbs -ej tbs 5 2

And passed *ms* the following combinations of parameters:

0.2999 0.95 0.2999
0.2999 0.8 0.2999
0.2999 0.65 0.2999
0.2999 0.5 0.2999
0.2999 0.35 0.2999
0.2999 0.2 0.2999

0.2999 0.05 0.2999
0.25 0.95 0.25
0.25 0.8 0.25
0.25 0.65 0.25
0.25 0.5 0.25
0.25 0.35 0.25
0.25 0.2 0.25
0.25 0.05 0.25
0.2 0.95 0.2
0.2 0.8 0.2
0.2 0.65 0.2
0.2 0.5 0.2
0.2 0.35 0.2
0.2 0.2 0.2
0.2 0.05 0.2
0.15 0.95 0.15
0.15 0.8 0.15
0.15 0.65 0.15
0.15 0.5 0.15
0.15 0.35 0.15
0.15 0.2 0.15
0.15 0.05 0.15
0.1 0.95 0.1
0.1 0.8 0.1
0.1 0.65 0.1
0.1 0.5 0.1
0.1 0.35 0.1
0.1 0.2 0.1
0.1 0.05 0.1
0.05 0.95 0.05
0.05 0.8 0.05
0.05 0.65 0.05
0.05 0.5 0.05
0.05 0.35 0.05
0.05 0.2 0.05
0.05 0.05 0.05
0.0001 0.95 0.0001
0.0001 0.8 0.0001
0.0001 0.65 0.0001
0.0001 0.5 0.0001
0.0001 0.35 0.0001
0.0001 0.2 0.0001
0.0001 0.05 0.0001

### $D_2$ Simulations

Our approach for simulating $D_2$ were largely similar to those used for $D_1$ except with different parameters, and the inclusion of simulations for the $B \to C$ direction of introgression in addition to $C \to B$. Like for $D_1$, we performed a small set of simulations to investigate introgression in both directions and verify our simulation approaches. For the method where parent trees were simulated separately and then combined, we used:

For parent tree 1:

ms 4 1000 -T -I 4 1 1 1 1 -ej 4.0 2 1 -ej 0.6 3 2 -ej 0.3 4 3
ms 4 1000 -T -I 4 1 1 1 1 -ej 4.0 2 1 -ej 0.65 3 2 -ej 0.3 4 3
ms 4 1000 -T -I 4 1 1 1 1 -ej 4.0 2 1 -ej 0.7 3 2 -ej 0.3 4 3
ms 4 1000 -T -I 4 1 1 1 1 -ej 4.0 2 1 -ej 0.75 3 2 -ej 0.3 4 3
ms 4 1000 -T -I 4 1 1 1 1 -ej 4.0 2 1 -ej 0.8 3 2 -ej 0.3 4 3
ms 4 1000 -T -I 4 1 1 1 1 -ej 4.0 2 1 -ej 0.85 3 2 -ej 0.3 4 3
ms 4 1000 -T -I 4 1 1 1 1 -ej 4.0 2 1 -ej 0.9 3 2 -ej 0.3 4 3

For parent tree 2:

ms 4 1000 -T -I 4 1 1 1 1 -ej 4.0 2 1 -ej 0.6 4 2 -ej 0.15 3 2
ms 4 1000 -T -I 4 1 1 1 1 -ej 4.0 2 1 -ej 0.65 4 2 -ej 0.15 3 2
ms 4 1000 -T -I 4 1 1 1 1 -ej 4.0 2 1 -ej 0.7 4 2 -ej 0.15 3 2
ms 4 1000 -T -I 4 1 1 1 1 -ej 4.0 2 1 -ej 0.75 4 2 -ej 0.15 3 2
ms 4 1000 -T -I 4 1 1 1 1 -ej 4.0 2 1 -ej 0.8 4 2 -ej 0.15 3 2
ms 4 1000 -T -I 4 1 1 1 1 -ej 4.0 2 1 -ej 0.85 4 2 -ej 0.15 3 2
ms 4 1000 -T -I 4 1 1 1 1 -ej 4.0 2 1 -ej 0.9 4 2 -ej 0.15 3 2

For parent tree 3:

ms 4 1000 -T -I 4 1 1 1 1 -ej 4.0 2 1 -ej 0.3 4 2 -ej 0.15 3 2

For the population split/rejoin method, we simulated the $C \to B$ and $B \to C$ directions of introgression only. This was done using the following commands:

For $C \to B$ introgression:

ms 4 2000 -T -I 4 1 1 1 1 -ej 4.0 2 1 -ej 0.6 3 2 -ej 0.3 4 3 -es 0.15 3 0.5 -ej 0.15 5 2
ms 4 2000 -T -I 4 1 1 1 1 -ej 4.0 2 1 -ej 0.65 3 2 -ej 0.3 4 3 -es 0.15 3 0.5 -ej 0.15 5 2
ms 4 2000 -T -I 4 1 1 1 1 -ej 4.0 2 1 -ej 0.7 3 2 -ej 0.3 4 3 -es 0.15 3 0.5 -ej 0.15 5 2
ms 4 2000 -T -I 4 1 1 1 1 -ej 4.0 2 1 -ej 0.75 3 2 -ej 0.3 4 3 -es 0.15 3 0.5 -ej 0.15 5 2
ms 4 2000 -T -I 4 1 1 1 1 -ej 4.0 2 1 -ej 0.8 3 2 -ej 0.3 4 3 -es 0.15 3 0.5 -ej 0.15 5 2
ms 4 2000 -T -I 4 1 1 1 1 -ej 4.0 2 1 -ej 0.85 3 2 -ej 0.3 4 3 -es 0.15 3 0.5 -ej 0.15 5 2
ms 4 2000 -T -I 4 1 1 1 1 -ej 4.0 2 1 -ej 0.9 3 2 -ej 0.3 4 3 -es 0.15 3 0.5 -ej 0.15 5 2

For $B \to C$ introgression:

ms 4 2000 -T -I 4 1 1 1 1 -ej 4.0 2 1 -ej 0.6 3 2 -ej 0.3 4 3 -es 0.15 2 0.5 -ej 0.15 5 3
ms 4 2000 -T -I 4 1 1 1 1 -ej 4.0 2 1 -ej 0.65 3 2 -ej 0.3 4 3 -es 0.15 2 0.5 -ej 0.15 5 3
ms 4 2000 -T -I 4 1 1 1 1 -ej 4.0 2 1 -ej 0.7 3 2 -ej 0.3 4 3 -es 0.15 2 0.5 -ej 0.15 5 3
ms 4 2000 -T -I 4 1 1 1 1 -ej 4.0 2 1 -ej 0.75 3 2 -ej 0.3 4 3 -es 0.15 2 0.5 -ej 0.15 5 3
ms 4 2000 -T -I 4 1 1 1 1 -ej 4.0 2 1 -ej 0.8 3 2 -ej 0.3 4 3 -es 0.15 2 0.5 -ej 0.15 5 3
ms 4 2000 -T -I 4 1 1 1 1 -ej 4.0 2 1 -ej 0.85 3 2 -ej 0.3 4 3 -es 0.15 2 0.5 -ej 0.15 5 3
ms 4 2000 -T -I 4 1 1 1 1 -ej 4.0 2 1 -ej 0.9 3 2 -ej 0.3 4 3 -es 0.15 2 0.5 -ej 0.15 5 3

#### *Simulating values of* $N_2$ *and* $N_3$

We used a similar approach to $D_1$ to simulate across different values of $N$. For $D_2$ we varied different splitting times and also did simulations in both directions of introgression. Parent tree 1 was combined with either parent tree

2 or 3 (but not both). $\gamma$ was held to 0.5 for all simulations varying $N$. For parent tree 1, we used the following command:

ms 4 49000 -t 0.01 -T -I 4 1 1 1 1 -ej 4.0 2 1 -ej tbs 3 2 -ej 0.2999 4 3

And passed the following parameters: 0.6, 0.65, 0.7, 0.75, 0.8, 0.85, 0.9. For parent tree 2, we used the following command:

ms 4 49000 -t 0.01 -T -I 4 1 1 1 1 -ej 4.0 2 1 -ej tbs 4 2 -ej 0.15 3 2 -en 0.15 2 tbs -en tbs 2 1

And passed the following combinations of parameters:

0.6 2 0.6
0.6 1.5 0.6
0.6 1.25 0.6
0.6 1 0.6
0.6 0.75 0.6
0.6 0.66 0.6
0.6 0.5 0.6
0.65 2 0.65
0.65 1.5 0.65
0.65 1.25 0.65
0.65 1 0.65
0.65 0.75 0.65
0.65 0.66 0.65
0.65 0.5 0.65
0.7 2 0.7
0.7 1.5 0.7
0.7 1.25 0.7
0.7 1 0.7
0.7 0.75 0.7
0.7 0.66 0.7
0.7 0.5 0.7
0.75 2 0.75
0.75 1.5 0.75
0.75 1.25 0.75
0.75 1 0.75
0.75 0.75 0.75
0.75 0.66 0.75
0.75 0.5 0.75
0.8 2 0.8
0.8 1.5 0.8
0.8 1.25 0.8
0.8 1 0.8
0.8 0.75 0.8
0.8 0.66 0.8
0.8 0.5 0.8
0.85 2 0.85
0.85 1.5 0.85
0.85 1.25 0.85
0.85 1 0.85
0.85 0.75 0.85
0.85 0.66 0.85

0.85 0.5 0.85
0.9 2 0.9
0.9 1.5 0.9
0.9 1.25 0.9
0.9 1 0.9
0.9 0.75 0.9
0.9 0.66 0.9
0.9 0.5 0.9

Lastly, we simulated parent tree 3 using the command:

ms 4 49000 -t 0.01 -T -I 4 1 1 1 1 -ej 4.0 2 1 -ej 0.3 4 2 -ej 0.15 3 2 -en 0.15 2 tbs -en 0.3 2 1

And passed it the following parameters: 2, 1.5, 1.25, 1, 0.75, 0.66, 0.5

#### *Simulating values of $\gamma_2$ and $\gamma_3$*

Like for $D_1$, we simulated across different admixture proportions using the population split/rejoin method, with a constant $N$ across the tree. For the $C \rightarrow B$ direction, we used the following command:

ms 4 98000 -t 0.01 -T -I 4 1 1 1 1 -ej 4.0 2 1 -ej tbs 3 2 -ej 0.3 4 3 -es 0.15 3 tbs -ej 0.15 5 2

And for the $B \rightarrow C$ direction, we used:

ms 4 98000 -t 0.01 -T -I 4 1 1 1 1 -ej 4.0 2 1 -ej tbs 3 2 -ej 0.3 4 3 -es 0.15 2 tbs -ej 0.15 5 3

Both of the above commands were passed the following combinations of parameters:

0.6 0.95
0.6 0.8
0.6 0.65
0.6 0.5
0.6 0.35
0.6 0.2
0.6 0.05
0.65 0.95
0.65 0.8
0.65 0.65
0.65 0.5
0.65 0.35
0.65 0.2
0.65 0.05
0.7 0.95
0.7 0.8
0.7 0.65
0.7 0.5
0.7 0.35
0.7 0.2
0.7 0.05
0.75 0.95
0.75 0.8
0.75 0.65

0.75 0.5
0.75 0.35
0.75 0.2
0.75 0.05
0.8 0.95
0.8 0.8
0.8 0.65
0.8 0.5
0.8 0.35
0.8 0.2
0.8 0.05
0.85 0.95
0.85 0.8
0.85 0.65
0.85 0.5
0.85 0.35
0.85 0.2
0.85 0.05
0.9 0.95
0.9 0.8
0.9 0.65
0.9 0.5
0.9 0.35
0.9 0.2
0.9 0.05

### Saccharomyces paradoxus *analysis*

#### *Estimation of demographic parameters*

For our analysis of the *Saccharomyces paradoxus* system, it was necessary for us to estimate several demographic parameters. First, we estimated the population-scaled mutation rate, $\theta$, using the genome wide average per-site heterozygosity, for the *SpB* and *SpC* populations. This was done using the following formulas from Chapter 3 of Hahn (2018):

$$\pi = \frac{\sum_{j=1}^{S} h_j}{G} \tag{32}$$

Where $G$ is the size of the genome in base pairs, $S$ is the number of segregating sites, and $h_j$ is the heterozygosity at segregating site $j$, defined for a single site as:

$$h = \frac{n}{n-1}(1 - \sum p_i^2) \tag{33}$$

Where $n$ is the number of sample sequences, and $p_i$ is the frequency of allele $i$ at the site. We used a genome size of $G = 1.2 * 10^7$ for this calculation. From this, we used the relationship $\theta = 4N\mu$, with a per-base mutation rate of $\mu = 1.84 * 10^{-10}$ (Fay & Benavides 2005, Zhu et al. 2014) to estimate approximate internal-branch effective population sizes for parent trees 1 and 2 of the system.

We estimated approximate lineage-splitting times in years for the system as $1.8 * 10^5$ for the outgroup, $t_2 = 1.0 * 10^5$, and $t_1 = 8.0 * 10^3$, based on supplementary figure S9 of Leducq *et al.* 2016. We took the mean value of our two population size estimates, $N = 6.5 * 10^5$, as the mean population size for the tree. Using this information, in combination with a generation time estimate of 2920 (Fay & Benavides 2005) generations per year, we were able to estimate our lineage-splitting times in units of 4*N* generations.

#### Simulating a hybrid speciation scenario

Using the above demographic parameter estimates, we simulated a scenario of hybrid speciation for the *S. paradoxus* system by assuming that the timing of migration $t_m$ equals $t_1$. Due to the lack of ILS in this system, we were able to simulate gene trees directly in the correct numbers; therefore, for each replicate simulation, we simulated 2002 trees for parent tree 1, corresponding to the ANC-topology windows, and 55 trees for parent tree 2, corresponding to the H0/H1b-topology windows. It was also not necessary to combine the parent trees into a single file; since we know the exact gene trees, we can compare them directly, as was done for the empirical estimate.

Since *ms* takes population size changes using a fold-difference, we set the whole tree equal to $\pi$ for the *SpC* population and then specified a 3.3-fold increase in the *SpB* population, corresponding to our observed difference. With these considerations in mind, we used this command to simulate the ANC-topology windows:

ms 4 2002 -t 0.000223 -T -I 4 1 1 1 1 -ej 202 2 1 -ej 112 3 2 -ej 20.2 4 3 -en 0 2 3.3

And this command to simulate the H0/H1b-topology windows:

ms 4 55 -t 0.000223 -T -I 4 1 1 1 1 -ej 202 2 1 -ej 112 4 2 -ej 20.2 3 2 -en 0 2 3.3

Simulated outputs from *ms* are in units of 4*N* generations, so we used our above estimates of $N$ and $\mu$ to convert them into percent divergence for comparison with our empirical estimate of $D_1$.

### Supplementary figures and tables

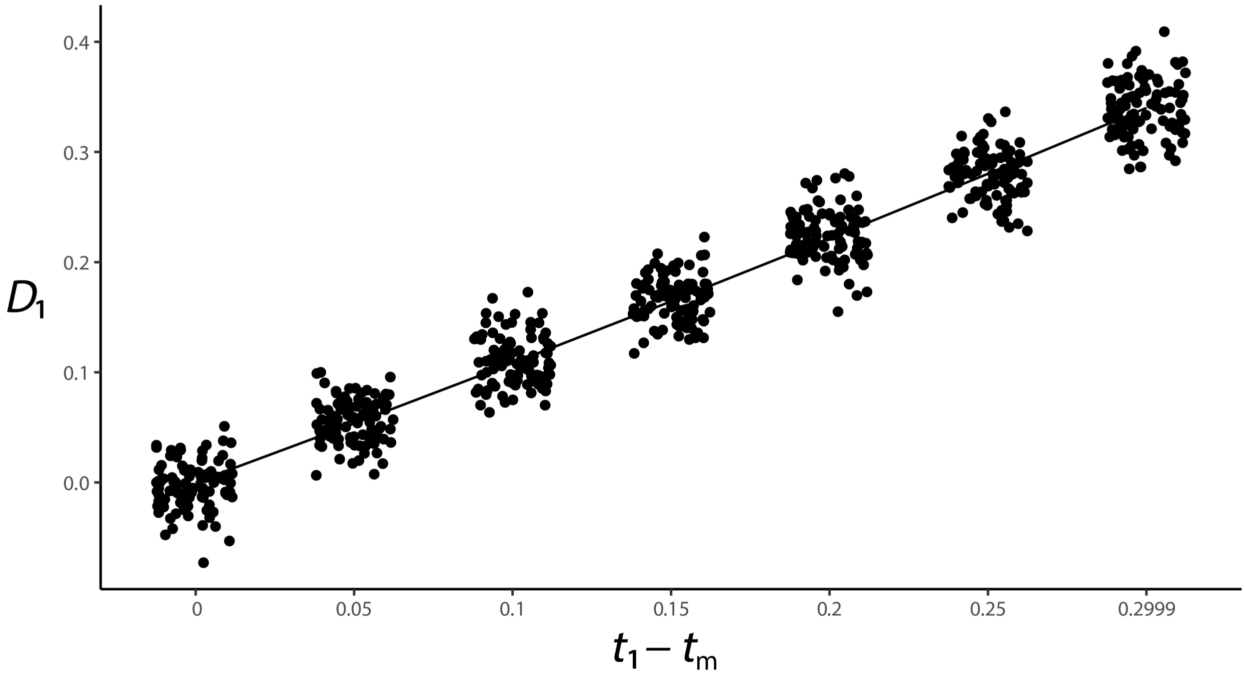


Figure S1: $D_1$ as a function of the difference in timing of speciation and introgression (units of $4N$), using an instantaneous population split and rejoin event to simulate introgression. Solid line indicates expected values from the mathematical model, while dots indicate simulated values at a particular timestep.

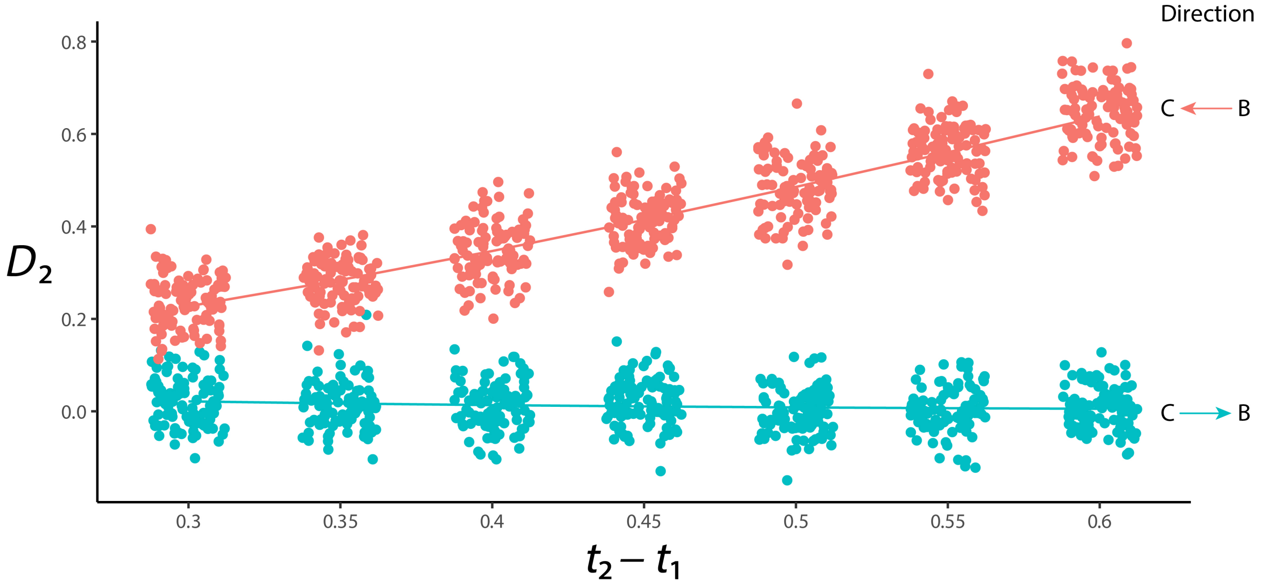


Figure S2: $D_2$ as a function of the time between speciation events in units of $4N$, using an instantaneous population split and rejoin event to simulate introgression. Color indicates the direction of introgression, solid line indicates mathematical expectation, and dots indicate simulated values.

| N1:N2 (t1 - tm = 0) | | | | | | |
|---|---|---|---|---|---|---|
| 0.5 | 0.66 | 0.8 | 1 | 1.33 | 1.5 | 2 |
| 0.9 | 0.18 | 0 | 0 | 0.13 | 0.35 | 0.92 |
| γ2 (t1 - tm = 0) | | | | | | |
| 0.05 | 0.2 | 0.35 | 0.5 | 0.65 | 0.8 | 0.95 |
| 1 | 1 | 0.85 | 0 | 0.98 | 1 | 1 |

Table S1: False positive rates for $D_1$. For each parameter value, this shows probability of incorrectly rejecting hybrid speciation based on the simulated null distribution ($t_1 - t_m = 0$, $N_1 : N_2 = 1$, $\gamma_2 = 0.5$) out of 100 simulated values.

| | | t1 - tm | | | | | |
|---|---|---|---|---|---|---|---|
| | | 0.05 | 0.1 | 0.15 | 0.2 | 0.25 | 0.2999 |
| N1:N2 | 0.5 | 0.85 | 1 | 0.85 | 0.15 | 0 | 0 |
| | 0.66 | 1 | 0.82 | 0.08 | 0 | 0 | 0 |
| | 0.8 | 1 | 0.34 | 0 | 0 | 0 | 0 |
| | 1 | 0.7 | 0 | 0 | 0 | 0 | 0 |
| | 1.33 | 0.04 | 0 | 0 | 0 | 0 | 0 |
| | 1.5 | 0.02 | 0 | 0 | 0 | 0 | 0 |
| | 2 | 0 | 0 | 0 | 0 | 0 | 0 |
| γ2 | 0.05 | 0 | 0 | 0 | 0 | 0 | 0 |
| | 0.2 | 0 | 0.01 | 0.21 | 0.69 | 0.99 | 0.91 |
| | 0.35 | 0.85 | 0.99 | 0.71 | 0.08 | 0 | 0 |
| | 0.5 | 0.58 | 0.01 | 0 | 0 | 0 | 0 |
| | 0.65 | 0 | 0 | 0 | 0 | 0 | 0 |
| | 0.8 | 0 | 0 | 0 | 0 | 0 | 0 |
| | 0.95 | 0 | 0 | 0 | 0 | 0 | 0 |

Table S2: False negative rates for $D_1$. For each combination of parameters, the probability of incorrectly accepting hybrid speciation based on the simulated null distribution ($t_1 - t_m = 0$, $N_1 : N_2 = 1$, $\gamma_2 = 0.5$) out of 100 simulated values.

| | | t2 - t1 | | | | | | |
|---|---|---|---|---|---|---|---|---|
| | | **0.3** | **0.35** | **0.4** | **0.45** | **0.5** | **0.55** | **0.6** |
| N1:N2 | 0.5 | 0.08 | 0.16 | 0.08 | 0.11 | 0.07 | 0.11 | 0.09 |
| | 0.66 | 0.08 | 0.01 | 0.09 | 0.11 | 0.06 | 0.06 | 0.03 |
| | 0.8 | 0.04 | 0.02 | 0.04 | 0.06 | 0.1 | 0.1 | 0.05 |
| | 1 | 0.06 | 0.04 | 0.04 | 0.02 | 0.06 | 0.03 | 0.06 |
| | 1.33 | 0.08 | 0.05 | 0.04 | 0.03 | 0.03 | 0.05 | 0.01 |
| | 1.5 | 0.05 | 0.08 | 0.03 | 0.03 | 0.02 | 0 | 0.03 |
| | 2 | 0.06 | 0.06 | 0.09 | 0.02 | 0.05 | 0.05 | 0.03 |
| γ2 | 0.05 | 0.98 | 0.96 | 0.97 | 0.96 | 0.93 | 0.95 | 0.97 |
| | 0.2 | 0.47 | 0.54 | 0.44 | 0.5 | 0.49 | 0.41 | 0.47 |
| | 0.35 | 0.12 | 0.11 | 0.17 | 0.12 | 0.14 | 0.14 | 0.09 |
| | 0.5 | 0.08 | 0.06 | 0.05 | 0.02 | 0.07 | 0.07 | 0.07 |
| | 0.65 | 0.3 | 0.28 | 0.14 | 0.1 | 0.06 | 0.09 | 0.06 |
| | 0.8 | 0.63 | 0.59 | 0.61 | 0.44 | 0.41 | 0.37 | 0.22 |
| | 0.95 | 0.97 | 0.97 | 0.95 | 0.96 | 0.96 | 0.87 | 0.91 |

Table S3: False positive rates for $D_2$. For each combination of parameters, the probability of incorrectly rejecting the $C \to B$ direction of introgression based on the simulated null distribution ($C \to B$ introgression, $N_1 : N_2 = 1$, $\gamma_2 = 0.5$) out of 100 simulated values.

| | | t2 - t1 | | | | | | |
|---|---|---|---|---|---|---|---|---|
| | | **0.3** | **0.35** | **0.4** | **0.45** | **0.5** | **0.55** | **0.6** |
| N1:N3 | 0.5 | 0.34 | 0.06 | 0.01 | 0 | 0 | 0 | 0 |
| | 0.66 | 0.12 | 0.01 | 0 | 0 | 0 | 0 | 0 |
| | 0.8 | 0.04 | 0.01 | 0 | 0 | 0 | 0 | 0 |
| | 1 | 0.04 | 0 | 0 | 0 | 0 | 0 | 0 |
| | 1.33 | 0 | 0 | 0 | 0 | 0 | 0 | 0 |
| | 1.5 | 0 | 0 | 0 | 0 | 0 | 0 | 0 |
| | 2 | 0 | 0 | 0 | 0 | 0 | 0 | 0 |
| γ3 | 0.05 | 0.19 | 0.21 | 0.34 | 0.35 | 0.6 | 0.6 | 0.72 |
| | 0.2 | 0.83 | 0.62 | 0.3 | 0.05 | 0 | 0 | 0 |
| | 0.35 | 0.19 | 0 | 0 | 0 | 0 | 0 | 0 |
| | 0.5 | 0.01 | 0 | 0 | 0 | 0 | 0 | 0 |
| | 0.65 | 0 | 0 | 0 | 0 | 0 | 0 | 0 |
| | 0.8 | 0 | 0 | 0 | 0 | 0 | 0 | 0 |
| | 0.95 | 0.06 | 0.06 | 0.02 | 0.05 | 0 | 0 | 0 |

Table S4: False negative rates for $D_2$. For each combination of parameters, the probability of incorrectly accepting the $C \rightarrow B$ direction of introgression based on the simulated null distribution ($C \rightarrow B$ introgression, $N_1 : N_2 = 1$, $\gamma_2 = 0.5$) out of 100 simulated values.
